# Plant miRNAs and amino acids interact to shape soil bacterial communities

**DOI:** 10.1101/2025.04.16.649127

**Authors:** Jessica A. Dozois, Marc-Antoine Duchesne, Katel Hallaf, Julien Tremblay, Étienne Yergeau

**Affiliations:** Institut National de la Recherche Scientifique, Centre Armand-Frappier Santé Biotechnologie, Laval, Québec, Canada

**Keywords:** plant miRNAs, amino acids, nitrogen, soil bacterial communities, plant-microbe interactions

## Abstract

Plants and microbes use many strategies to acquire soil amino acids. Recent findings suggest that genes related to amino acid metabolism and transport are influenced by plant miRNAs. This was the first report of a plant-derived molecule with the potential to modify microbial amino acid uptake. Here, we first show that *Arabidopsis* modifies its root miRNA content when fertilized with a mixture of 17 amino acids. The miRNAs that responded to amino acid fertilization and other rhizosphere-abundant miRNAs were applied to a simplified soil community, grown with diverse amino acid sources, to test if they interfered with microbial community growth, community composition and amino acid consumption. Plant miRNAs affected the community’s growth in over 70 % of the amino acid sources. The effectiveness of plant miRNAs also depended on the N source supplied to the microbial community, with the strongest effect observed with L-lysine. Specifically, ath-miR159a reduced the microbial consumption of L-lysine, further supporting that plant miRNAs can influence microbial amino acid uptake. Plant miRNAs also affected the relative abundance of specific bacterial taxa, which we subsequently isolated. All isolates were affected in terms of growth when exposed to miRNAs. Two out of three isolates were also impacted in their amino acid consumption. Surprisingly, while plant miRNAs inhibited amino acid consumption at both the community and isolate levels, they had mostly positive effects. Our results suggest that rhizospheric plant miRNAs might have a role in modulating the amino acid consumption of soil bacteria, but not necessarily in a competitive framework.

## Introduction

Nitrogen (N) is one of the most limiting nutrients in terrestrial ecosystems – hence the subject of a fierce competition between plants and microorganisms [1]. Recent findings from our team revealed that plant miRNAs impact the expression of many bacterial genes related to N, including genes related to amino acid uptake and metabolism [2]. Although the concentration of amino acids in the soil is rather low (∼20 µM) [3], the pool is rapidly renewed within minutes to hours [4–6]. The uptake of amino acids by microbes exceeds, by a factor >8, the rate at which they convert organic N to ammonium and nitrate [7]. When amino acid concentrations are high, plants uptake less nitrate [8–10]. Amino acids are thus sought after by microbes and plants. To impair plant amino acid uptake and increase the efflux of amino acids from the roots, microbes such as *Fusarium* can produce zearalenone while *Pseudomonas*, *Chromobacterium* can produce 2,4-diacetylphloroglucinol and isolates belonging to the phyla *Actinomycetota* and *Pseudomonadota* can produce phenazine [11–13]. To better colonize the root-soil interface, cytokinin-producing bacteria, like *Bacillus subtilis*, can also increase amino acid rhizodeposition [14]. Although microbes uptake soil amino acids more rapidly, plants have high affinity transporters for amino acids [15] and are competitive in environmental conditions such as soil acidification [16]. Plants also stimulate microbes to depolymerize soil organic nitrogen and in time uptake N released from the microbial necromass [17]. Some plants directly influence soil N-cycling by producing compounds that inhibit microbes from performing nitrification and denitrification [18–24]. However, if confirmed, plant miRNAs would be the first molecule that plants use to interfere with microbial amino acid uptake. Within the plant, specific miRNAs regulate N metabolism and respond to different exogenous N treatments [25–31], but their effect on the microbial community is not known.

The effectiveness of RNA interference (RNAi) across kingdoms has been shown for two miRNAs in cotton plants (*Gossypium*) that inhibited the virulence of a fungal pathogen (*Verticillium dahliae*) [32]. Plant miRNAs and small RNAs are also exchanged from plants to animals [33–39] where they have been found to affect host cell activity and regulate the gut microbiota. This RNA cross-talk has also been shown from plants to plants [40, 41], from plants to phytopathogens [32, 42], from microbial pathogens to plants [43, 44] and from plant symbionts to plants [45]. Our team recently discovered that plant miRNAs are found in the rhizosphere and that they affect microbial communities [2]. Transcriptomic analysis of the key rhizosphere bacterium *Variovorax paradoxus* [46] exposed to plant miRNAs revealed hundreds of differentially expressed genes [2]. Since, as mentioned above, most of the highly affected genes were in the COG category “amino acid metabolism and transportation”, we sought to further confirm the link between bacteria, amino acids and plant miRNAs. We conducted three independent experiments to test the hypotheses that 1. exposing a soil microbial community to plant miRNAs in various amino acid sources will impact microbial community growth and amino acid uptake, 2. that specific bacteria will respond to miRNAs leading to shifts in the microbial community composition and 3. that plant miRNAs will reduce the amino acid consumption of the miRNA-responsive bacteria.

We first looked at the miRNAs in *Arabidopsis* roots treated with amino acids or ammonium nitrate. The miRNAs that responded positively to amino acids were mixed with those commonly found in the rhizosphere and used in vitro with a simplified soil community cultured with various amino acid sources. Bacteria that responded to miRNAs in some amino acid sources were then isolated and individually confronted to the miRNAs to assess their impact on growth and amino acid consumption.

## Material and methods

A more descriptive version of the methods is available in the supplementary material.

### Profiles of miRNA and the bacterial community in response to fertilizer treatments *in planta*

#### Experimental design

*Arabidopsis thalian*a Col-0 (n=5) were grown under three different nitrogen treatments that were supplied every 2-3 days: a mix of 17 L-amino acids (0.190 g of N/L), a no added nitrogen control and an inorganic nitrogen control (ammonium nitrate, 0.190 g of N/L). These controls enabled us to differentiate the influence of 1) nitrogen rich fertilizers vs. no added nitrogen and 2) amino acids vs. an inorganic N source on the relative abundance of miRNAs and on the bacterial community. The growth chamber was set to 18 hours of daylight and 6 hours of darkness. After 21 days, the roots, rhizosphere, and distant soil were sampled and flash-frozen in liquid nitrogen.

#### miRNA profiling

The miRNA profiles were obtained by small RNA sequencing on RNA extracted from roots (RNeasy Plant Mini, Qiagen) (Illumina HiSeq4000, Centre d’expertise et de services de Génome Québec, Montreal, Canada). To associate small RNA reads to specific miRNAs [2, 47], the reads were pre-processed and trimmed [48], filtered for quality and common contaminants were removed (bbduk). These reads were mapped against a reference genome: *A. thaliana* (TAIR10/GCA_000001735.1). Alignment was carried out using the BWA parameters aln mismatch=1 and seed=5. Finally, the potential miRNAs that mapped against the genome were compared (BLASTn) to the miRBase hairpin and miRBase mature databases. To accurately identify the plant miRNAs induced by our fertilizer treatments, we removed all miRNAs that had been identified in our unplanted soil controls.

#### Bacterial community profiling

Since plant-microbe interactions are central to our hypotheses, we characterized the bacterial community of the *in planta* experiment via amplicon sequencing of the V4-V5 region of the 16S rRNA gene (515F-Y and 926R [49]) on a MiSeq apparatus (Illumina) at the National Research Council of Canada, (Montreal, Canada) from RNA extracted from roots. For microbial taxonomic labelling, we treated amplicon sequencing data with the pipeline AmpliconTagger [50]. This pipeline grouped the sequences into *amplicon sequence variants* (ASVs) (100% identical sequences) [51] and identified their taxonomic identity with RDP classifier using the SILVA R138 database [52–54].

#### Correlations and linear models linking miRNAs and bacteria

To link given taxa and our miRNAs of interest, Spearman correlations and linear regression models were performed on their relative abundances in the roots of *A. thaliana*. We tested the assumptions of linearity, homoscedasticity and normality before running the regression models.

### Effects of synthetic miRNAs on a microbial community grown with different amino acids

#### Soil microbial community

We developed a simplified soil community by adding 2 g of sieved agricultural soil from our experimental field (45.5416°N, 73.7173°W) in five growth media. After 28 hours (200 rpm, 25 °C), the cultures were filtered (30 µm), normalized to the same optical density, then pelleted (4 °C, 15 min, 4700 *g*). The pellets were suspended in PBS, pooled, and aliquoted into sterile cryotubes in a cryoprotective solution [55] before being stored at −80 °C.

#### Microbial growth

To verify whether miRNAs interfered with microbial activity depending on the amino acid source, five were tested. The miRNAs were synthesized with the 3’ end 2’-OH methylation specific to plant miRNAs (Integrated DNA Technologies, Supplementary Table 1) [56, 57]. The effect of these five synthetic plant miRNAs (2 µM for each miRNA) on the activity of soil microbes was evaluated. As a control, scrambled miRNAs corresponding to the five plant miRNAs were used (i.e., the sequences of ribo nucleic acids were randomized). To test if plant miRNAs modified microbial activity, soil microbes were grown in media containing a mixture of all 17 L-amino acids (15 mM) or in media containing individual amino acids (15 mM). In all media, artificial root exudates (glucose, fructose, sucrose, lactic acid, succinic acid and citric acid) [58] served as an additional carbon source (15 mM). We added a tetrazolium dye which turns purple in the presence of active dehydrogenases and enhance optical density. The optical density (600 nm) was measured every hour with a plate reader (Tecan Infinite M1000Pro). When microbial activity was different between the miRNA treatment and scrambled control, we repeated the experiment with individual miRNAs (2 µM) to identify which of the five miRNAs was responsible for the change.

#### Amino acid quantification

We quantified the effect of individual miRNA (2 µM) on the microbial consumption of four different amino acid sources: L-proline, L-lysine, glycine, and the mix of 17 AA. The microbial community was sampled at the beginning of the experiment, at the early log-phase, the mid-log phase and the stationary phase. The amino acids were quantified (in technical duplicates) using their respective standard curves in a colorimetric assay. For quantifying L-proline and L-lysine, we added our samples to a reaction mix consisting of 1% ninhydrin, 60% acetic acid, and 20% ethanol [59] and incubated the mixture at 95°C for 20 min. Whereas for glycine, the reaction mix consisted of 1% ninhydrin diluted in 80% ethanol and was incubated for 15 min at 75°C. The optical density was read at 520 nm for L-proline and glycine and 540 nm for L-lysine (Tecan Infinite M1000Pro). The mix of 17 AA was quantified with the colorimetric L-Amino Acid Assay Kit (CELL BIOLABS INC. MET-5054) as specified by the supplier, except for the standard curve which was our own 17-AA mix. The optical density measurements were converted to amino acid concentration (µM) using the standard curve.

#### Bacterial community composition

We then evaluated if the change in growth was reflected in a change within the bacterial community. We sampled the community at the endpoint of the experiment (52 h) and DNA was extracted with a physical microvolume extraction [60] and purified via ethanol-NaCl precipitation. The libraries were prepared and the V4-V5 region of the 16S rRNA gene 515F-Y and 926R was sequenced [49] (Illumina MiSeq, Centre d’expertise et de services de Génome Québec, Montreal, Canada). We labelled the taxa with RDP classifier and SILVA [52–54] as previously described. We identified specific ASVs for which the relative abundance significantly changed in response to the miRNA treatment according to both the DESeq2 [61] and ANCOM-BC [62] analyses.

### Isolates challenged with miRNAs

#### Isolation of strains from the simplified soil community

We isolated the bacteria by using different solid media including a Phosphate Separately autoclaved Reasoner’s 2A meant to isolate *Chryseobacterium* and *Flavobacterium* [63]. We extracted the DNA of the isolates with a physical microvolume extraction [60], amplified the V4-V5 16S rRNA gene region, purified the PCR products (QIAquick PCR Purification Kit, Qiagen) and sent the samples for Sanger sequencing (Centre d’expertise et de services de Génome Québec, Montreal, Canada) (forward primer: 515FY and reverse primer:926R [49]). Consensus sequences were generated with the BioEdit Sequence Alignment Editor and compared to the 16S rRNA sequences of our previously identified ASVs with BLASTn. The isolates that had perfect matches with the responsive ASVs were further used.

#### Isolates of interest exposed to miRNAs

We cultured three of our isolates (*Acinetobacter*, *Chryseobacterium* and *Raoultella*) overnight in rich media, washed the cells, normalized them (DO600=2.00) and challenged them to 2 µM of individual miRNAs or 10 µM of the mix as well as the corresponding scrambled controls. The isolates were grown in media, previously described, consisting of 15 mM of amino acids and 15 mM of artificial root exudates. To stay coherent with the community experiment, we added the tetrazolium dye and measured the optical density (OD600) every hour for 52 hours. We repeated the experiment five times and used paired T-tests to find timepoints and growth phases where optical density and the area under the curve differed statistically between the miRNA plant and scrambled treatments.

To quantify the effect of miRNAs on the amino acid uptake of each isolate, the isolates were grown in the 17 AA mix medium and sampled at four different timepoints. The amino acids were quantified using the colorimetric L-Amino Acid Assay Kit (CELL BIOLABS INC. MET-5054) as previously described. We then calculated if the isolate’s growth and amino acid use differed between the miRNA plant and scrambled treatments. Paired T-tests were used for each time point and for the area under the curve, provided the assumptions were met. Otherwise, the Wilcoxon paired test was used.

#### Whole Genome sequencing of isolates

Isolates were cultured overnight (28 °C, 200 rpm) in in minimal medium and DNA was extracted with QIAmp DNA Mini Kit (Qiagen) following the protocol for Gram negative bacteria. Quantity and quality of DNA were verified prior to library preparation and Nanopore sequencing (PromethION, Oxford Nanopore technologies). The reads were combined per isolate and filtered for quality with filtlong (https://github.com/rrwick/Filtlong). The assembly was performed with Flye [64] and the expected genome size for each genus. The assembled genomes were annotated with NCBI Prokaryotic Genome Annotation Pipeline (PGAP) (https://www.ncbi.nlm.nih.gov/refseq/annotation_prok/).

#### Putative miRNA targets within the assembled genomes

Since the functional pairing between plant miRNAs and bacterial mRNA has yet to be validated, we used three tools to predict plant miRNA bacterial targets: psRNATarget [65], miRanda [66, 67] and BLASTn for short queries. We only considered targets that were common between the three tools. The sequences of all 10 miRNAs (five plant sequences and five scrambled sequences) were mapped to the coding DNA sequences (CDS) of each isolate that are known to be involved in amino acid transport or general nitrogen regulation.

#### Co-culture of Arabidopsis thaliana and isolates

To test the relationship between *A. thaliana* and each of the three isolates, a gnotobiotic assay was performed with surface sterilized seeds. On the day of the experiment, the seeds were inoculated with washed cultures of each isolate for 2 hours. The optical density at 600 nm (OD600) for each isolate was previously calculated to correspond to a concentration of 10^4^ CFU/µl. The negative control culture medium, which was washed alongside the cultures, was used to inoculate the seeds. Ten seeds were then placed in a sterile Magenta Box containing half-strength MS (Murashige and Skoog) medium with 0.8 % agar. The growth chamber settings were the same as in the previous *Arabidopsis* growth experiment. The experiment lasted for 23 days. We then determined if isolates impacted *Arabidopsis* growth by comparing different plant traits with Kruskal-Wallis tests for each time point and for the area under the curve.

## Results

### Root miRNAs shift in response to amino acids

Amino acids increased the relative abundance of six root miRNAs (ath-miR158b, ath-miR160a-3p, ath-miR166b-5p, ath-miR390a-3p, ath-miR827 and ath-miR5642b) (Supplementary Fig.1). Among the 10 most abundant root miRNAs, the relative abundance of only ath-miR408 was different among N treatments (Supplementary Fig. 2): this miRNA was less abundant under amino acid fertilization than the NH_4_NO_3_ treatment. Five miRNA candidates were selected for further experiments: miRNAs that positively responded to the amino acid treatment (ath-miR158b, ath-miR827 and ath-miR5642b) and miRNAs abundant in the rhizosphere that were shown to shift the amino acid gene expression in *Variovorax paradoxus* (ath-miR158a-3p and ath-miR159a) [2] (Fig. 1).

**Figure 1:**
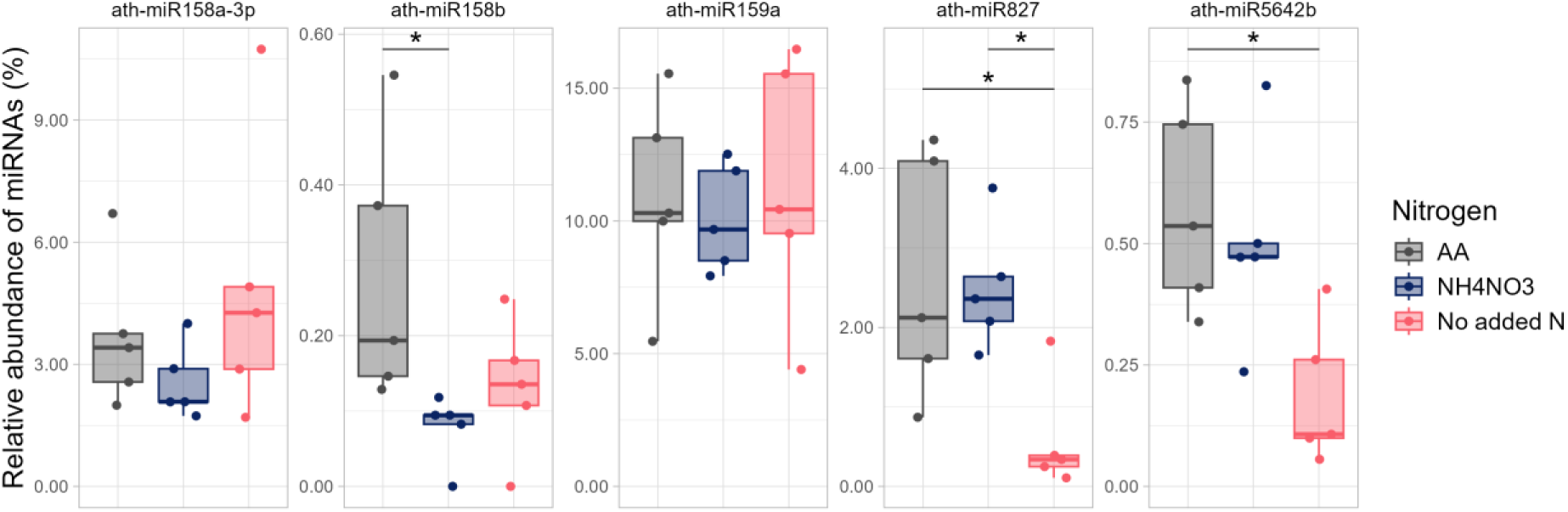
The relative abundance of the five selected miRNAs inside the roots of *A. thaliana* depending on the nitrogen treatment. (AA: a mix of amino acids, NH_4_NO_3_: Ammonium nitrate and No added N). Differences in relative abundance of miRNAs between N treatments are indicated with brackets (*p*-adjusted with Holm correction <0.05).

### Bacterial ASVs correlate to the relative abundance of miRNAs

To highlight links between the relative abundance of certain taxa and the relative abundance of miRNAs, we performed Spearman correlations and compiled those that were significant (*p*<0.05) (Supplementary Table 2). Although there were no correlations for ath-miR158b, three were significant for ath-miR158a-3p, one for ath-miR159a, five for ath-miR827 and three for ath-miR5642b (Supplementary Fig. 3). Half of the correlations were positive, and the others were negative. Among the taxa correlated to miRNA, some also responded to our N treatments (Supplementary Fig. 4). Five of our correlations involved N-responding miRNAs and N-responding taxa, suggesting that indirect miRNA-microbe interactions were more likely at play. With linear regression models, we confirmed that the N treatments were also important in influencing the abundance of *Massilia* (ASV#14) and *Niastella* (ASV#67) (Supplementary Table 3). When the variable “N treatments” was partialled out from our linear regression models, 7/12 of the ASV-miRNA pairs were still significant (*p*<0.05, Table 1). The models that were more suggestive of direct miRNA-microbe interactions included: Ktedonobacteria (ASV#111)∼ ath-miR158a-3p, Ktedonobacteria (ASV#111)∼ath-miR827 and *Litorilituus* (ASV#630)∼ath-miR158a-3p.

**Table 1:**
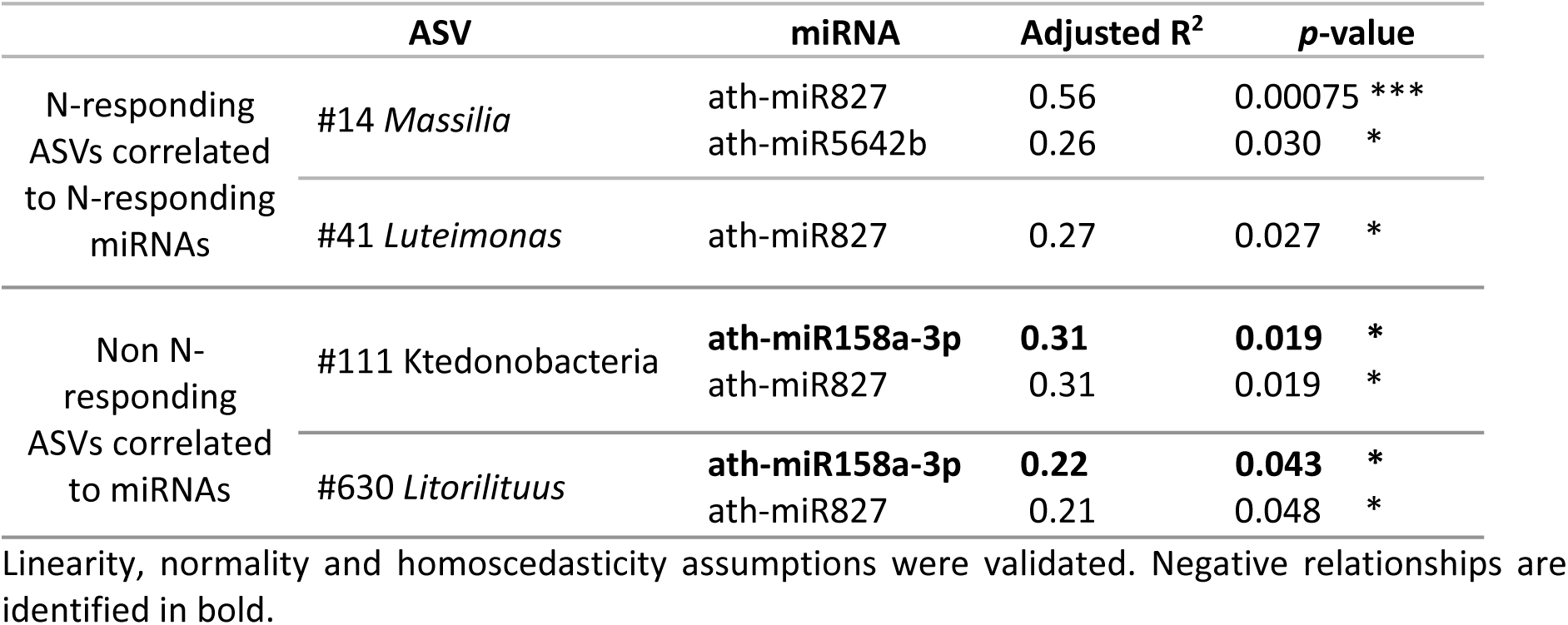
Linear regressions between the relative abundances of miRNA and bacterial taxa.

### Plant miRNAs impact the growth of soil microbes and the amino acid use of bacterial isolates

We first evaluated if the five miRNAs (Fig. 1 and Supplementary Table 1) interfered with microbial growth in different amino acid sources. The mixture of miRNAs increased or decreased the growth of the community for at least two consecutive timepoints (2 hours) for 13 out of the 18 amino acid sources (Table 2 and Supplementary Fig. 5). For most of the amino acid sources, the effect of miRNAs on microbial growth occurred during the exponential phase (Fig. 2 and Supplementary Table 4). To assess miRNA effects on community growth, two nitrogen sources were tested for negative impacts (L-proline, glycine) and two for positive impacts (17 L-amino acid mix, L-lysine). For L-proline and glycine, ath-miR827 and ath-miR5642b reproduced the negative effect (Supplementary Fig. 6, Supplementary Table 5). For the mix of 17 L-AA and L-lysine both ath-miR159a and ath-miR827 had positive effects. ath-miR158a-3p only positively affected microbial growth in L-lysine (Supplementary Fig. 6, Supplementary Table 5). However, the effects that lasted throughout the exponential growth phase were all positive (Supplementary Table 5). The bacterial community responded more strongly to miRNAs when cultured in L-lysine (responded to 3/5 miRNAs) where ath-miR159a produced the most lasting effects (Supplementary Table 5). For each of the miRNA-AA source pairs found in Supplementary Fig. 6, the effect of miRNAs on microbial AA consumption was also tested (Supplementary Fig. 7). Microbes reduced their consumption of L-lysine when exposed to ath-miR159a (2 µM) compared to its scrambled control (Fig. 3A) whereas miRNAs did not impact the uptake of any other amino acids.

**Figure 2:**
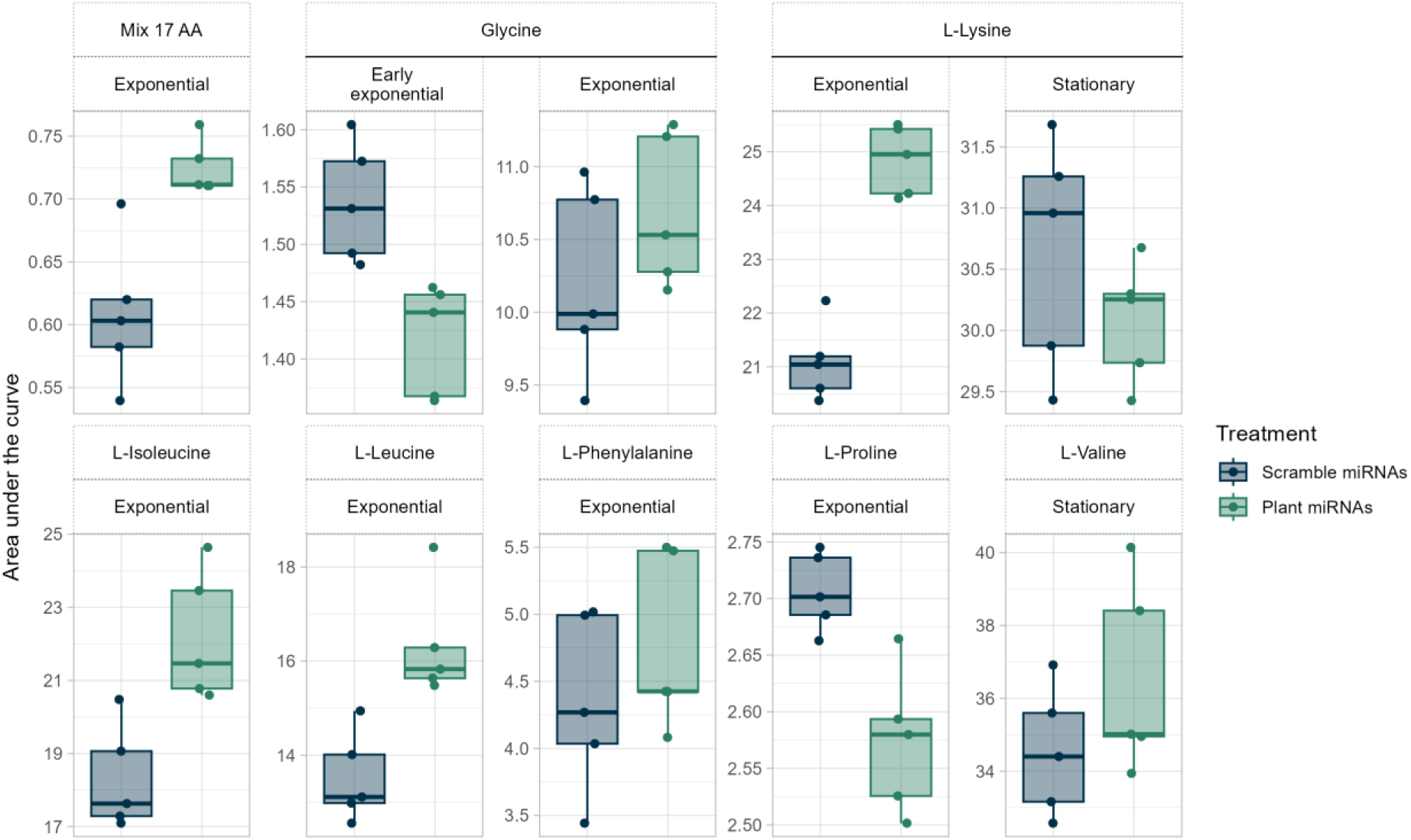
The miRNA treatments impact the area under the growth curve during key growth phases. The simplified soil community’s growth phases were different (*p* <0.05, paired T-test) between the miRNA treatments (10 µM of mix of plant miRNAs vs. 10 µM of a mix of scrambled miRNAs) in these eight amino acid sources.

**Figure 3.**
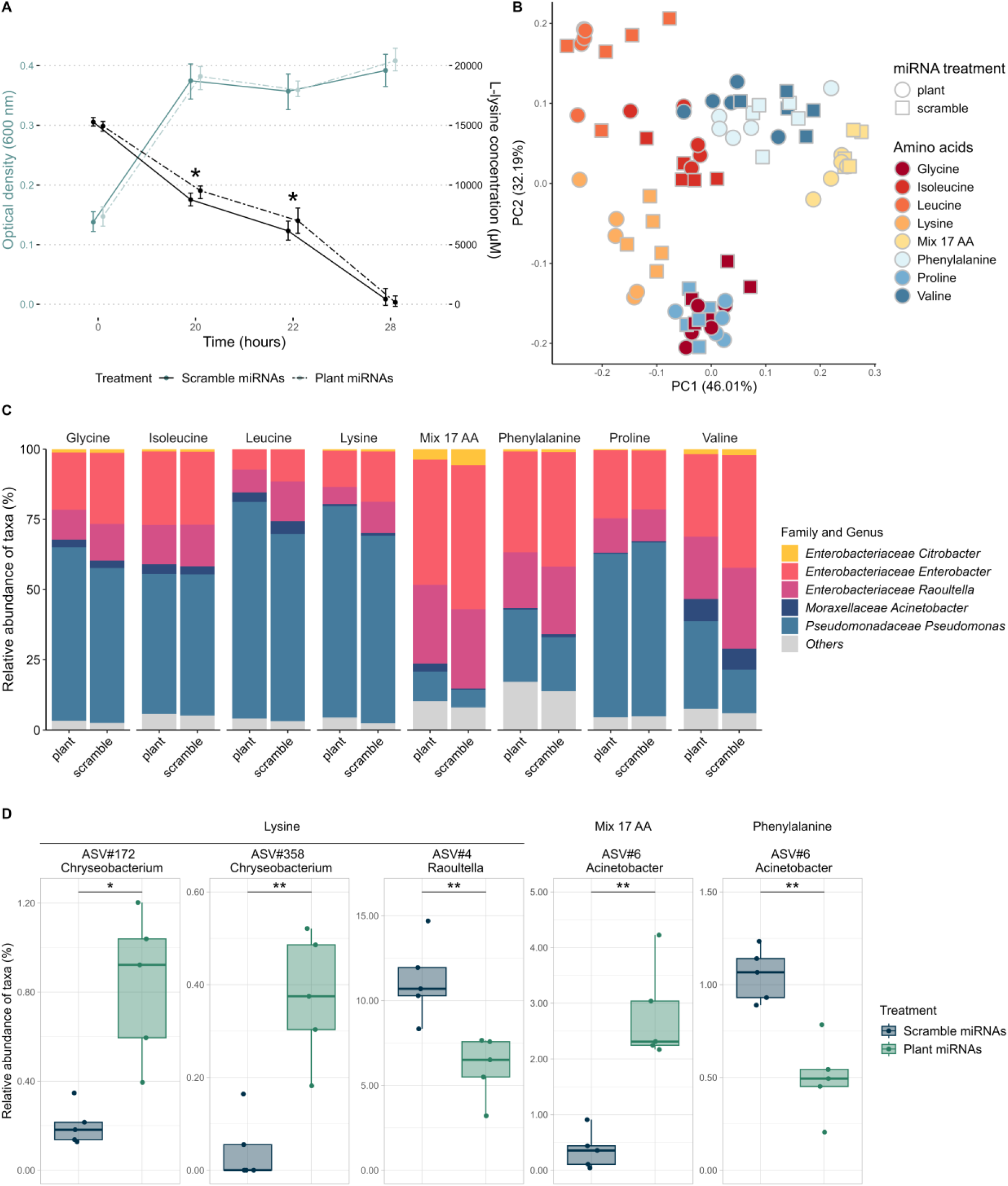
miRNAs modify bacterial composition and L-lysine consumption. A. Microbial growth (OD600, blue lines) and L-lysine consumption (black lines) over time (hours). Dashed lines indicate that the microbes were exposed to 2 µM of ath-miR159a whereas full lines indicate that the microbes were exposed to 2 µM of scrambled ath-miR159a. Significant results are identified with a * (*p*<0.05 paired T-test). B. Principal component analysis of bacteria grown in different amino acid sources and exposed to the mix of miRNAs (10 µM). C. Composition of the soil microbial community grown with different amino acids as a nitrogen source. D. ASVs for which the relative abundance was different (*p* <0.05) between the miRNA treatments (10 µM of mix of plant miRNAs vs. 10 µM of a mix of scrambled miRNAs). The relative abundance of the ASVs was found to be significantly different for both DESeq2 and ANCOM-BC analyses.

**Table 2:**
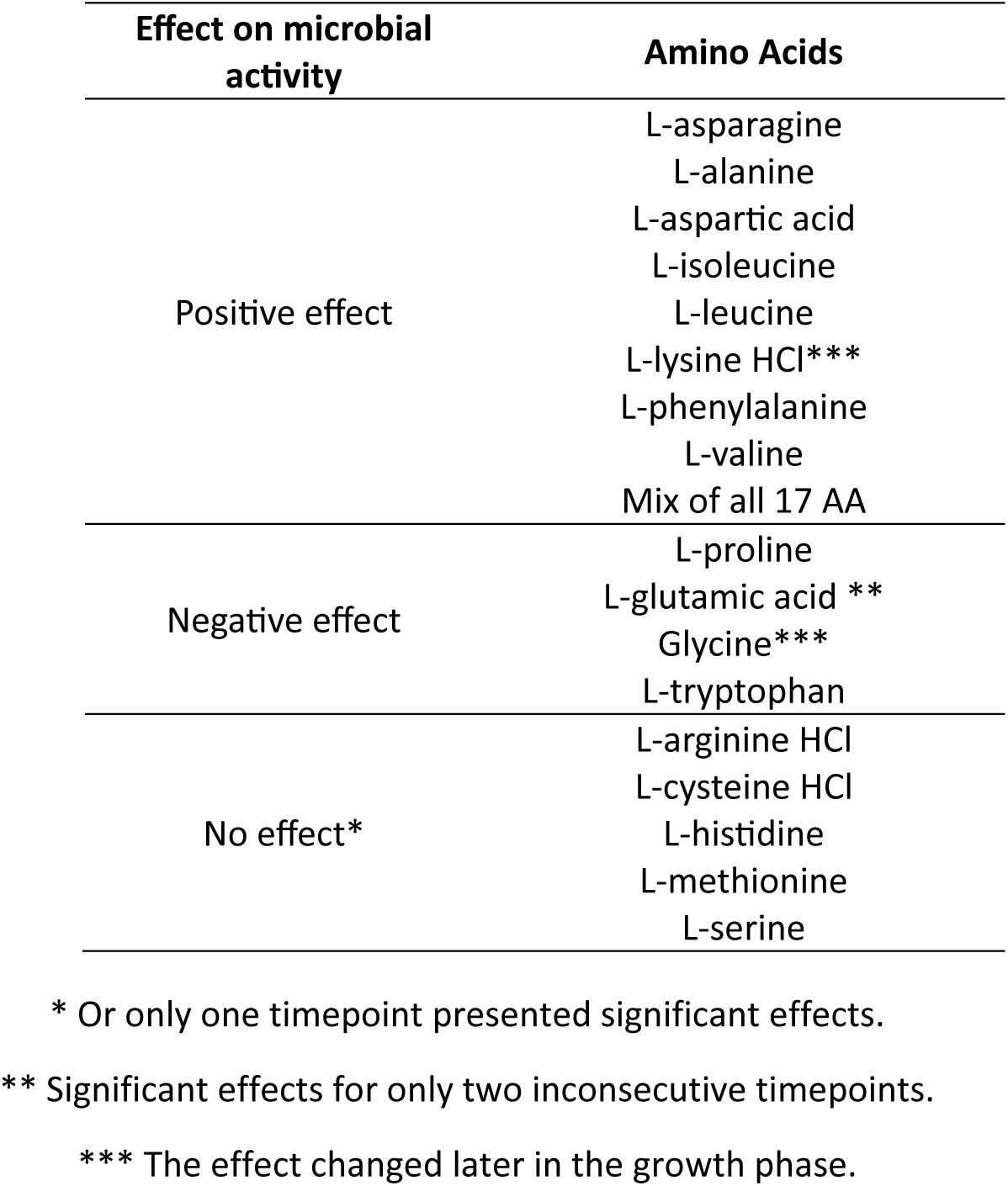
Effect of 10 µM of miRNAs (plant miRNAs compared to scrambled miRNAs) on the activity (OD600) of soil microbes.

Taxonomic bacterial changes caused by miRNAs were investigated for eight amino acids (glycine, L-isoleucine, L-leucine, L-lysine, L-phenylalanine, L-proline, L-valine, Mix of 17 AA) by 16S rRNA gene sequencing (Fig. 3B and 3C). Our medium, composed of 17 amino acids, favored bacteria of the family Enterobacteriaceae compared to media containing one type of amino acid. The bacterial community exposed to a single miRNA was mainly influenced by the type of amino acid provided (R^2^ = 0.1427, *p*=0.001, Supplementary Table 6, Supplementary Fig. 8) whereas the bacterial community exposed to a mix of miRNAs was influenced by the interaction of the miRNA treatment (plant vs. scrambled) and the type of amino acid provided (R^2^ =0.1478, *p*=0.013, Supplementary Table 6).

Microbial community changes, such as a lower abundance of *Raoultella* and *Enterobacter,* were induced by the mix of plant miRNAs at the family and genus level for most nitrogen sources (*p*<0.05). Interestingly, L-isoleucine instigated no obvious microbial changes at this taxonomic level (Fig. 3C). At the ASV level, we identified four ASVs differentially abundant between the miRNA treatments (plant vs. scrambled) (*p*<0.05): ASV#4-*Raoultella,* ASV#6-*Acinetobacter*, ASV#172-*Chryseobacterium* and ASV#358-*Chryseobacterium* (Fig. 3D). Plant miRNAs negatively impacted the relative abundance of *Raoultella* (ASV#4) grown in L-lysine. The other taxa *Chryseobacterium* (ASV#172 and ASV#358) cultured in L-lysine were positively affected by plant miRNAs. A particularity occurred with ASV#6-*Acinetobacter* which seemed to prosper when exposed to plant miRNAs in a mix of amino acids but exhibited the opposite phenotype when cultured in L-phenylalanine.

The response of the isolates corresponding to ASV#4-*Raoultella,* ASV#6-*Acinetobacter*, ASV#172-*Chryseobacterium* was confirmed. The largest changes in isolate growth were caused by ath-miR158b (Supplementary Tables 7 and 8). The positive effects of plant miRNAs on the growth of *Acinetobacter* cultured in the AA mix were coherent with the effects at the community level (Fig. 3D). Though, the amino acid uptake of *Acinetobacter* was only higher in response to the mix of plant miRNAs at the endpoint of the experiment (Fig. 4A). When *Acinetobacte*r was grown in L-phenylalanine, the mix of miRNAs only positively impacted the growth which was unexpected, though 48 h post-inoculation some of the replicates treated with the scrambled control had a growth spurt (Supplementary Fig. 9). Based on the effects at the community level, we also expected the mix of five plant miRNAs to positively impact *Chryseobacterium* which was reconfirmed in the amino acid consumption experiment (Fig. 4B). Yet, in isolation, ath-miR158b caused the opposite effect (Fig. 4A, Supplementary Tables 7 & 8 and Supplementary Fig.9). The prolonged negative effect of ath-miR158b on the growth of *Raoultella*, may also explain its negative response to plant miRNAs within the simplified microbial community. *Raoultella* was the only isolate both negatively impacted by the mix of five plant miRNAs in the soil community and in pure culture. The mix of five miRNAs and the plant miRNA ath-miR158b reduced the amino acid uptake of *Raoultella* (Fig. 4AB). Yet, we found that plant miRNAs ath-miR158a-3p and ath-miR5642b could stimulate higher amino acid uptake within a 24-hour period (Fig. 4B). Curiously, ath-miR5642b initially stimulated amino acid uptake at the start of the experiment, but three hours later, it delayed both amino acid uptake and growth compared to the scrambled control. Whole genome sequencing revealed that *Raoultella* had the most diverse set of inorganic nitrogen and amino acid transporters (Supplementary Table 9) suggesting its potential to mitigate the negative effects caused by a specific miRNA.

**Figure 4:**
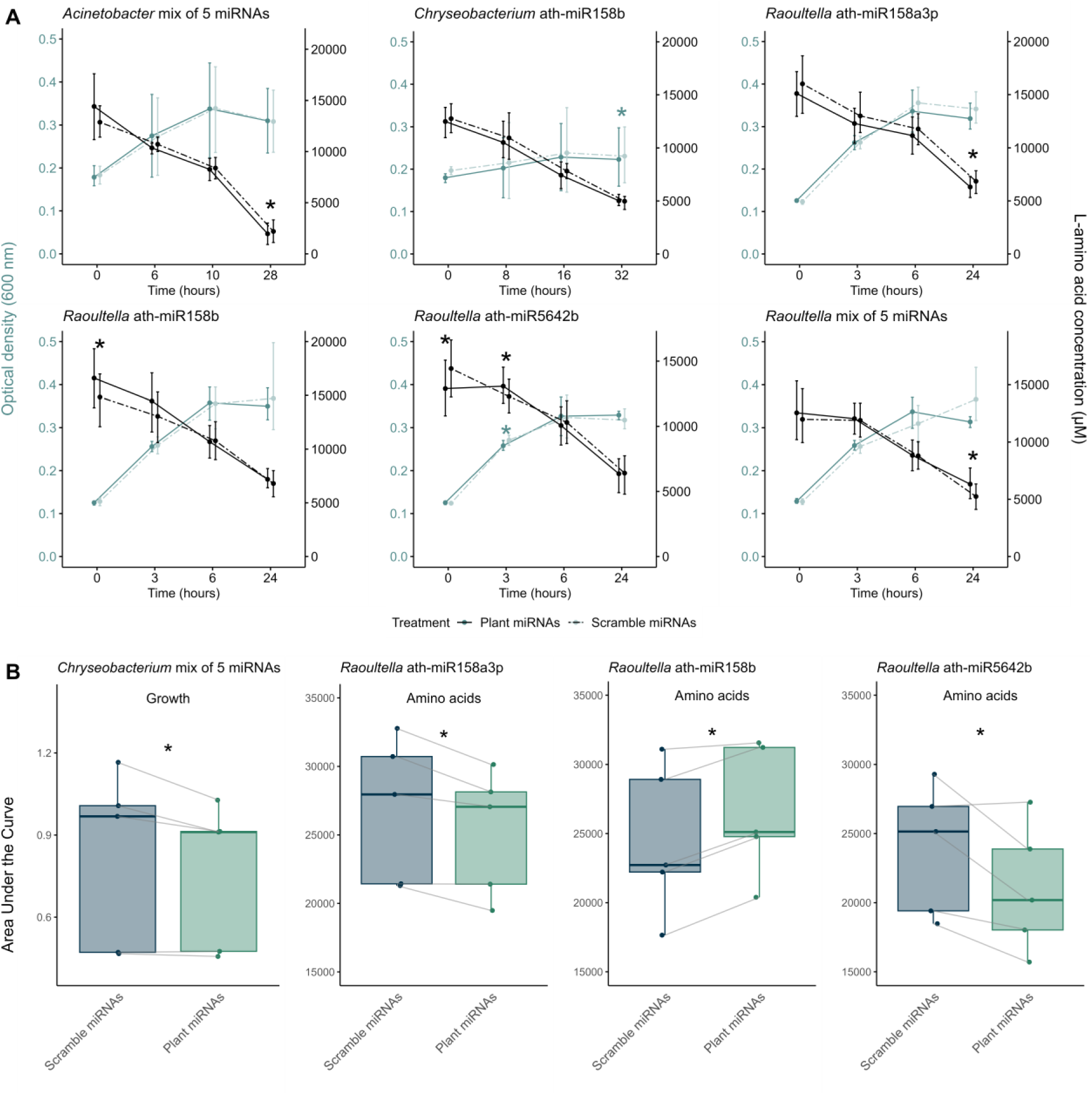
The miRNA treatments affect how isolates grow and the rate at which they uptake amino acids from the 17 AA mix media. A. Microbial growth (OD600, blue lines) and amino acid consumption (black lines) over time (hours). Full lines indicate that the microbes were exposed to plant miRNAs whereas dashed lines indicate that the microbes were exposed to scrambled miRNAs. Significant results are identified with a * (*p*<0.05 paired T-test) (n=5). B. Differences in the area under the curve for *Chryseobacterium* growth and for *Raoultella* amino acid uptake (*p*<0.05 paired T-test) (n=5).

### miRNAs ath-miR159a and ath-miR827 potentially target more amino acid transportation genes

Our target predictions identified a total of fifteen plant miRNA targets and six scrambled miRNA targets (Supplementary Table 10). A total of seven targets were identified for *Acinetobacter*, none for *Chryseobacterium* and fourteen targets for *Raoultella*. This was consistent with the number of coding sequences related to amino acid transportation and nitrogen regulation we identified for each isolate: 19 CDS, 4 CDS, 67 CDS for *Acinetobacter, Chryseobacterium* and *Raoultella* respectively. For *Acinetobacter*, ath-miR159a targeted the most genes including a sodium/proline symporter, a serine/threonine transporter and a glnG nitrogen regulation protein NR(I) while its scrambled equivalent targeted none. For *Raoultella,* ath-miR827 targeted the most genes with six different targets including: a high-affinity branched-chain amino acid ABC transporter permease, an abgT p-aminobenzoyl-glutamate transporter, a proline-specific permease and a few amino acid ABC transporters permeases/ATP-binding proteins. No targets were identified for the scrambled version of ath-miR827. For *Raoultella,* ath-miR5642b also targeted three different genes: a glycine betaine/L-proline transporter, an aromatic amino acid DMT transporter and a tryptophan permease whereas its scrambled version targeted none. Overall, the miRNAs that targeted the most genes related to amino acid transportation were ath-miR159a, for *Acinetobacter*, and ath-miR827 for *Raoultella*. These miRNAs were respectively found to have the greatest impact on the soil community’s amino acid use and growth

### The isolate *Raoultella* delayed the germination of *Arabidopsis* and in time killed the plants

After observing the different impacts of miRNAs on isolate growth and amino acid utilization, we explored whether these effects could be explained by how the isolates influence the germination and development of *Arabidopsi*s. The plants inoculated with *Raoultella* still managed to germinate (Fig. 5A), though they never developed a floral stem (Fig. 5B). Of the three isolates, only *Raoultella* delayed *Arabidopsis* germination and killed the plants (Fig. 5CD).

**Figure 5:**
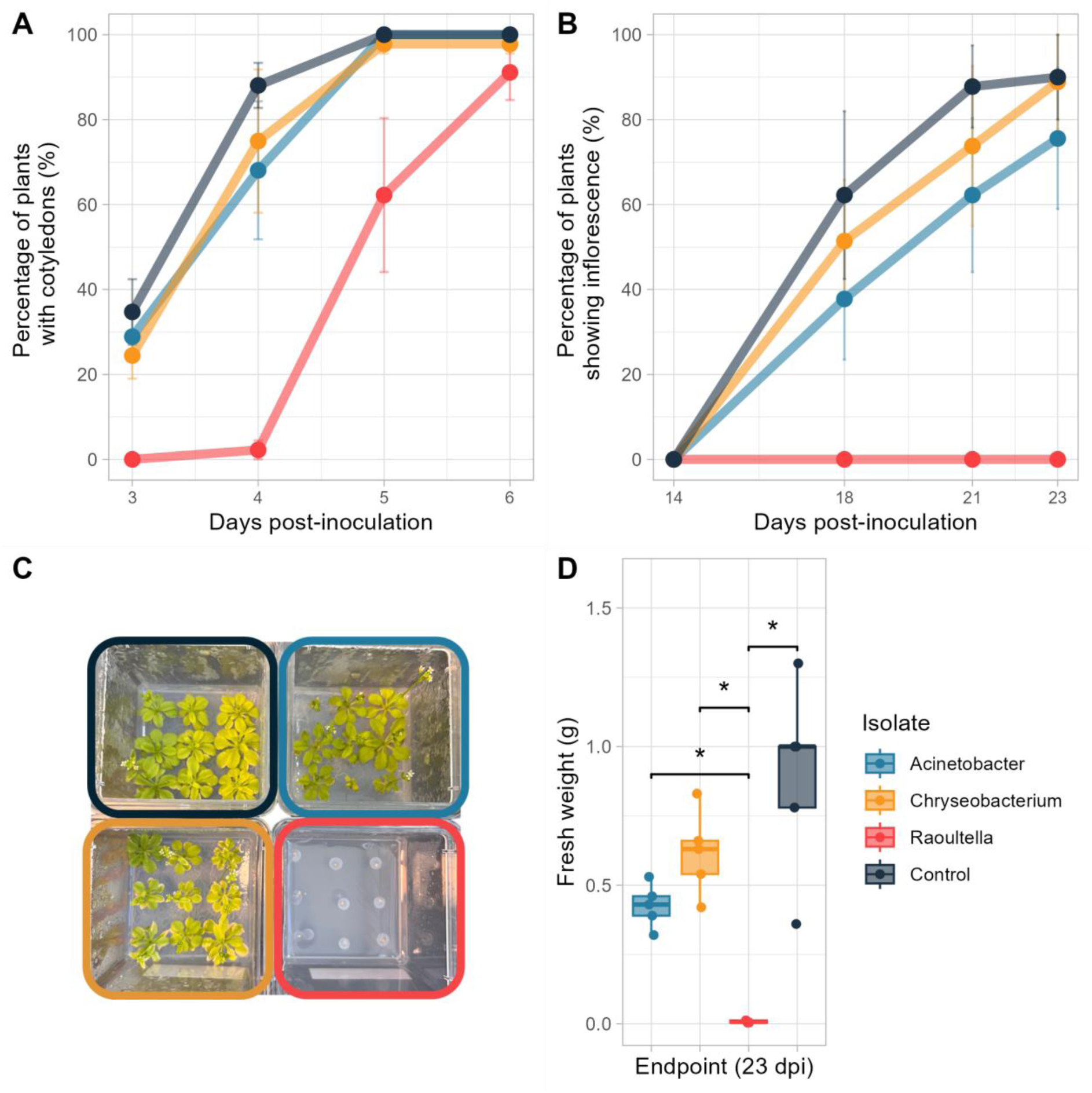
Effect of the three isolates (*Acinetobacter*: light blue, *Chryseobacterium*: yellow, *Raoultella*: red) on *Arabidopsis* germination and development. A. Percentage of plants with cotyledons, B. Percentage of plants showing inflorescence (floral stem), C. Picture of plants 23 days post-inoculation and D. Fresh weight of the plants 23 days post-inoculation. (n=5)

## Discussion

We had previously shown that *V. paradoxus* – a key rhizosphere bacterium – shifted the expression of many genes in the “amino acid transport and metabolism” COG category in response to exposure to plant miRNAs [2]. Here we show that 1. amino acid fertilization changes the relative abundance of miRNAs in *Arabidopsis* roots; 2. as hypothesized these miRNAs affect a simplified soil bacterial community, where ath-miR827 most effectively alters soil bacterial growth, 3. miRNAs elicit the strongest bacterial response when they are grown in L-lysine, and 4. plant miRNAs impact the growth and amino acid use of certain isolates from the simplified soil bacterial community. As expected, we detected miRNA-responsive bacteria, affected in terms of growth and amino acid uptake, within the soil community. However, the predominantly positive effects of plant miRNAs were unexpected.

We identified a dozen of miRNAs in the roots of *A. thaliana* that responded to amino acids, many of which were among the most abundant miRNAs in the rhizosphere [2]. Within the plant, specific miRNAs regulate N metabolism and respond to different exogenous N treatments [25–29]. For instance, ath-miR827 is known to contribute to phosphate homeostasis in a nitrate-dependant manner [28, 68–70]. Like previous research [28, 69], we also found ath-miR827 to be less abundant in the roots of *A. thaliana* in low N conditions. We found ath-miR827 to be the most correlated to bacterial taxa in plant roots and it was retained in 4 out of 7 of our regression models. However, in this *in planta experiment*, it was challenging to separate the direct effects of amino acids on the bacterial community from the indirect effects due to shifts in plant miRNAs. Indeed, amino acids can influence microbe-microbe interactions, community composition, and functional diversity by serving as carbon and nitrogen sources and precursors for bioactive molecules that promote plant growth and facilitate microbe signaling [71, 72]. Some amino acids, like L-tryptophan, can enhance bacterial functions, while others, such as L-methionine, L-valine, L-cysteine, and L-serine, can inhibit growth and IAA production [73]. Amino acids like L-phenylalanine can drive stronger bacterial shifts through bioactive molecule production or selective pressure [74]. To distinguish direct from indirect effects, we conducted *in vitro* experiments with a simplified soil bacterial community. We chose to test N-responding miRNAs (ath-miR158b, ath-miR827 and ath-miR5642b) and miRNAs known to be abundant in the rhizosphere (ath-miR158a-3p and ath-miR159a).

As previously reported, we showed here that bacterial community and isolates can respond in a contradictory manner to the presence of different plant miRNAs. The first case is when a single miRNA has a different effect on different bacteria. Previously, bol-miR159 triggered taxa-dependent negative (*Bacillus*) or positive (*Weissella* and *Ralstonia*) effects [39]. In another study, among four to five human miRNAs predicted to target two gut bacteria, only one was shown to promote the growth of *Fusobacterium nucleatum* (hsa-miR-515-5p) and one to promote the growth of *Escherichia coli* (hsa-miR-1226-5p) suggesting again that miRNAs differentially affect bacteria [38]. In a mixed community assessed using methods that produce proportional data – such as 16S rRNA gene sequencing –, this positive effect could be due to an inhibition of a community member that would result in an apparent increase in other community members. To clarify this, we also isolated three of the most responsive community members and confirmed that they also responded in isolation, both positively and negatively. During stationary growth, the mixture of five plant miRNAs promoted amino acid consumption by *Acinetobacter* while reducing it in *Raoultella*. All three isolates showed varying growth responses to plant miRNAs: *Acinetobacter* grew more, *Raoultella* grew less, and *Chryseobacterium* was affected either way depending on the miRNA. This suggests that – in contrast to our hypothesis – the effect of plant miRNAs extends beyond competitive interactions and may be involved in the enrichment of specific bacteria. Although most reports of sRNAs exchange involve a plant host and a pathogen, *Rhizobium* can deliver sRNA fragments to soybean cells to control nodule initiation and development [75]. Arbuscular mycorrhizal fungi, another key symbiont, have also been shown to exchange sRNAs with the plant to benefit their mutualistic interactions [45, 76, 77]. It appears that a miRNA could be used by the plant to fine-tune the microbial community composition, by either inhibiting or enhancing the activities of different members of the microbiota.

The second case is when different miRNAs have different effects on the same bacterium. For instance, the growth of *Chryseobacterium* in the mixture of amino acids was negatively impacted by ath-miR158b and ath-miR827, and positively by ath-miR5642b, whereas the growth of *Raoultella* in L-Lysine was positively affected by ath-miR159a and negatively by ath-miR158a-3p, ath-miR158b and ath-miR5642b. Similarly, different plant miRNAs were shown to induce different responses in growth and activity of *Lactobacillus*, meaning that a combination of miRNAs is unlikely to have synergetic effect [33]. Here, the community composition of our simplified soil bacterial community responded more to a combination of miRNAs than to single miRNAs. At a community level, a diverse mixture of miRNAs interacting with a range of bacterial targets would more likely result in a detectable effect. The variety of miRNAs likely interferes with different pathways within various bacterial species, leading to stronger responses. Here again, this suggests that the plant could use subtle variations in the miRNA content of its rhizosphere to fine-tune the microbial community composition.

On top of these miRNA- and taxa-specific responses of the bacterial community, the responses were shown here to vary with amino acids. For instance, the soil community growth was both positively (mix of 17AA and L-lysine) and negatively (glycine and L-proline) affected by ath-miR827. This miRNA has also been shown to limit the entry of *Lactobacillus rhamnosus* into gut epithelial cells [33], but we show here that its effect is modulated by nutrients. Similarly, nutrients have been shown to modulate DNA uptake: transformation of *Pseudomonas stutzeri* was more frequent at the onset of the stationary phase under nutrient limitation [78], while for *Acinetobacter calcoaceticus*, a nutrient upshift with phosphate salts in the soil microcosms enhanced transformation rates [79]. Also, microbes cultured with L-lysine were more responsive to plant miRNAs. L-lysine has been shown to increase cell permeability or cell wall damage of bacteria such as *Acinetobacter baumannii*, *Escherichia* coli, *Klebsiella pneumoniae*, making them more susceptible to antibiotics [80, 81]. Perhaps bacteria grown in L-lysine were also more likely to take up miRNAs because of an increased cell permeability. Lysine is a particularly interesting amino acid, as it may be protected from soil microbial decay because of its positive charge which allows it to bind to negatively charged soil particles [82, 83]. Plants have Lysine-Histidine Transporters (LHT) [84, 85] and Amino Acid Permeases (APP) [86, 87] that can uptake L-lysine from the soil making them good competitors for this slowly degrading N source. Although L-lysine is found in root exudates [88, 89] and the increased leaching of lysine could benefit pathogens, as L-lysine is a key precursor for plant systemic acquired resistance (SAR) to infections [90]. L-lysine has also been shown to serve as a nutrient source for the soil bacterium *Pseudomonas putida* [91, 92], suggesting that it also helps microbes colonize the rhizosphere. Since we showed that *Arabidopsis* modulates the miRNA composition of its root environment, depending on the amount and type of nitrogen available, we could also envision indirect – through miRNAs – effects of nutrient availability on the bacterial community, on top of the direct effects discussed here. In fact, in our *in planta* experiment we found some significant relationships between individual rhizosphere bacteria and plant miRNAs even when partialling out the effect of fertilization. This suggests that, through a nutrient-induced modulation of root miRNA, plants could influence the microbial community composition.

Among our microbial community, *Raoultella* appeared as the most affected by plant miRNAs. *Raoultella* was indeed the only isolate, both negatively impacted by the mix of five plant miRNAs in the simplified soil community and in pure culture. We hypothesized that *Raoultella* would be detrimental for plant growth and confirmed that it negatively impacted *Arabidopsis* germination and growth. Our isolate also had numerous amino acid transporters encoded in its genome, which is typical for highly competitive bacteria and/or pathogens [93–95]. Consequently, the plant miRNAs used were predicted to target more genes in the *Raoultella* genome as compared to the other isolates. This overrepresentation of target genes might explain why we found more coherent effects of the various miRNAs under different conditions for *Raoultella*, as compared to *Chryseobacterium* and *Acinetobacter*. These two latter bacteria were sometimes favored and sometimes inhibited by plant miRNAs, had lower amounts of N-related genes in their genomes – and consequently harbored less predicted targets –, and finally did not have detrimental effects on *Arabidopsis*. We speculate that the increased presence of N-related genes – a hallmark of pathogens and efficient competitors for N – could suffice for theses bacteria to be more targeted by plant miRNAs.

Here we showed that plant miRNAs modify bacterial growth, relative abundance, and amino acid consumption, depending on the amino acid source supplemented. Taken together our results suggest that plants use miRNAs to fine tune the microbial community depending on the soil nutrient status. Having miRNAs that can increase or decrease various community members depending on their competitive or cooperative nature under nutrient limitation or not would be immensely advantageous for plants. It also suggests an avenue to use miRNAs to modify plant-associated microbial communities, towards improving agricultural sustainability.

## Acknowledgements

We extend our gratitude to colleagues and conference attendees for engaging in passionate conversations about our work. We are thankful to the Platform for the Characterization of Biological and Synthetic Nanovehicles (VBS) of INRS and Frédéric Veyrier for Nanopore sequencing, to Alex Rivera Millot for DNA extractions and to Luke Harrison for bioinformatic analyses.

## Data availability statement

The sequencing data generated in this study has been deposited under NCBI BioProject accessions PRJNA1107220 (root miRNAs), PRJNA1248534 (root, rhizosphere and bulk soil 16S rRNA gene amplicons), PRJNA1111829 (simplified soil community 16S rRNA gene amplicons) and PRJNA1247481 (whole genome sequence of isolates). The R code used to analyse the data and generate the figures is available on GitHub (https://github.com/le-labo-yergeau/Dozois_AA_miRNAs).

## References

1. L’Espérance E, Bouyoucef LS, Dozois JA et al. Tipping the plant-microbe competition for nitrogen in agricultural soils. iScience. 2024;27:110973 10.1016/j.isci.2024.110973

2. Middleton H, Dozois JA, Monard C et al. Rhizospheric mirnas affect the plant microbiota. Isme Communications. 2024;4 10.1093/ismeco/ycae120

3. Jones DL, Kielland K, Sinclair FL et al. Soil organic nitrogen mineralization across a global latitudinal gradient. Global Biogeochemical Cycles. 2009;23 10.1029/2008GB003250

4. Glanville HC, Hill PW, Schnepf A et al. Combined use of empirical data and mathematical modelling to better estimate the microbial turnover of isotopically labelled carbon substrates in soil. Soil Biology and Biochemistry. 2016;94:154–68 10.1016/j.soilbio.2015.11.016

5. Jones DL, Kielland K. Soil amino acid turnover dominates the nitrogen flux in permafrost-dominated taiga forest soils. Soil Biology and Biochemistry. 2002;34:209–19 10.1016/S0038-0717(01)00175-4

6. Kielland K, McFarland JW, Ruess RW et al. Rapid cycling of organic nitrogen in taiga forest ecosystems. Ecosystems. 2007;10:360–68 10.1007/s10021-007-9037-8

7. Wanek W, Mooshammer M, Blöchl A et al. Determination of gross rates of amino acid production and immobilization in decomposing leaf litter by a novel 15n isotope pool dilution technique. Soil Biology and Biochemistry. 2010;42:1293–302 10.1016/j.soilbio.2010.04.001

8. Nazoa P, Vidmar JJ, Tranbarger TJ et al. Regulation of the nitrate transporter gene in responses to nitrate, amino acids and developmental stage. Plant Molecular Biology. 2003;52:689–703 Doi 10.1023/A:1024899808018

9. Gioseffi E, de Neergaard A, Schjoerring JK. Interactions between uptake of amino acids and inorganic nitrogen in wheat plants. Biogeosciences. 2012;9:1509–18 10.5194/bg-9-1509-2012

10. Vidmar JJ, Zhuo D, Siddiqi MY et al. Regulation of high-affinity nitrate transporter genes and high-affinity nitrate influx by nitrogen pools in roots of barley. Plant Physiol. 2000;123:307–18 10.1104/pp.123.1.307

11. Phillips DA, Fox TC, King MD et al. Microbial products trigger amino acid exudation from plant roots. Plant Physiol. 2004;136:2887–94 10.1104/pp.104.044222

12. Johnson ET, Bowman MJ, Gomes RP et al. Identification of 2,4-diacetylphloroglucinol production in the genus. Scientific Reports. 2023;13 10.1038/s41598-023-41277-0

13. Dar D, Thomashow LS, Weller DM et al. Global landscape of phenazine biosynthesis and biodegradation reveals species-specific colonization patterns in agricultural soils and crop microbiomes. Elife. 2020;9 10.7554/eLife.59726

14. Kudoyarova GR, Melentiev AI, Martynenko EV et al. Cytokinin producing bacteria stimulate amino acid deposition by wheat roots. Plant Physiol Biochem. 2014;83:285–91 10.1016/j.plaphy.2014.08.015

15. Fischer WN, André B, Rentsch D et al. Amino acid transport in plants. Trends in Plant Science. 1998;3:188–95 10.1016/S1360-1385(98)01231-X

16. Pan WK, Tang S, Zhou JJ et al. Plant-microbial competition for amino acids depends on soil acidity and the microbial community. Plant and Soil. 2022;475:457–71 10.1007/s11104-022-05381-w

17. Kuzyakov Y, Xu X. Competition between roots and microorganisms for nitrogen: Mechanisms and ecological relevance. New Phytol. 2013;198:656–69 10.1111/nph.12235

18. Subbarao GV, Ishikawa T, Ito O et al. A bioluminescence assay to detect nitrification inhibitors released from plant roots: A case study with brachiaria humidicola. Plant and Soil. 2006;288:101–12 10.1007/s11104-006-9094-3

19. Subbarao GV, Yoshihashi T, Worthington M et al. Suppression of soil nitrification by plants. Plant Sci. 2015;233:155–64 10.1016/j.plantsci.2015.01.012

20. Zakir HAKM, Subbarao GV, Pearse SJ et al. Detection, isolation and characterization of a root-exuded compound, methyl 3-(4-hydroxyphenyl) propionate, responsible for biological nitrification inhibition by sorghum (sorghum bicolor). New Phytologist. 2008;180:442–51 10.1111/j.1469-8137.2008.02576.x

21. Subbarao GV, Nakahara K, Hurtado MP et al. Evidence for biological nitrification inhibition in brachiaria pastures. Proc Natl Acad Sci U S A. 2009;106:17302–7 10.1073/pnas.0903694106

22. Subbarao GV, Nakahara K, Ishikawa T et al. Biological nitrification inhibition (bni) activity in sorghum and its characterization. Plant and Soil. 2013;366:243–59 10.1007/s11104-012-1419-9

23. Bardon C, Poly F, Piola F et al. Mechanism of biological denitrification inhibition: Procyanidins induce an allosteric transition of the membrane-bound nitrate reductase through membrane alteration. FEMS Microbiol Ecol. 2016;92:fiw034 10.1093/femsec/fiw034

24. Bardon C, Piola F, Bellvert F et al. Evidence for biological denitrification inhibition (bdi) by plant secondary metabolites. New Phytol. 2014;204:620–30 10.1111/nph.12944

25. Cai H, Lu Y, Xie W et al. Transcriptome response to nitrogen starvation in rice. Journal Biosciences. 2012;37:731–47 10.1007/s12038-012-9242-2

26. Nguyen GN, Rothstein SJ, Spangenberg G et al. Role of micrornas involved in plant response to nitrogen and phosphorous limiting conditions. Frontiers in Plant Science. 2015;6 10.3389/fpls.2015.00629

27. Sinha SK, Rani M, Bansal N et al. Nitrate starvation induced changes in root system architecture, carbon:Nitrogen metabolism, and mirna expression in nitrogen-responsive wheat genotypes. Appl Biochem Biotechnol. 2015;177:1299–312 10.1007/s12010-015-1815-8

28. Zhao M, Tai H, Sun S et al. Cloning and characterization of maize mirnas involved in responses to nitrogen deficiency. PLoS One. 2012;7:e29669 10.1371/journal.pone.0029669

29. Zuluaga DL, De Paola D, Janni M et al. Durum wheat mirnas in response to nitrogen starvation at the grain filling stage. PLoS One. 2017;12:e0183253 10.1371/journal.pone.0183253

30. Das S, Singh D, Meena HS et al. Long term nitrogen deficiency alters expression of mirnas and alters nitrogen metabolism and root architecture in indian dwarf wheat (triticum sphaerococcum perc.) genotypes. Sci Rep. 2023;13:5002 10.1038/s41598-023-31278-4

31. Li L, Li Q, Davis KE et al. Response of root growth and development to nitrogen and potassium deficiency as well as microrna-mediated mechanism in peanut (arachis hypogaea l.). Front Plant Sci. 2021;12:695234 10.3389/fpls.2021.695234

32. Zhang T, Zhao YL, Zhao JH et al. Cotton plants export micrornas to inhibit virulence gene expression in a fungal pathogen. Nat Plants. 2016;2:16153 10.1038/nplants.2016.153

33. Teng Y, Ren Y, Sayed M et al. Plant-derived exosomal micrornas shape the gut microbiota. Cell Host Microbe. 2018;24:637–52.e8 10.1016/j.chom.2018.10.001

34. Mu J, Zhuang X, Wang Q et al. Interspecies communication between plant and mouse gut host cells through edible plant derived exosome-like nanoparticles. Mol Nutr Food Res. 2014;58:1561–73 10.1002/mnfr.201300729

35. Vaucheret H, Chupeau Y. Ingested plant mirnas regulate gene expression in animals. Cell Res. 2012;22:3–5 10.1038/cr.2011.164

36. Zhang L, Hou D, Chen X et al. Exogenous plant mir168a specifically targets mammalian ldlrap1: Evidence of cross-kingdom regulation by microrna. Cell Res. 2012;22:107–26 10.1038/cr.2011.158

37. Li M, Chen T, Wang R et al. Plant mir156 regulates intestinal growth in mammals by targeting the wnt/β-catenin pathway. Am J Physiol Cell Physiol. 2019;317:C434–C48 10.1152/ajpcell.00030.2019

38. Liu S, da Cunha A, Rezende R et al. The host shapes the gut microbiota via fecal microrna. Cell Host Microbe. 2016;19:32–43 10.1016/j.chom.2015.12.005

39. Xu Q, Qin XS, Zhang Y et al. Plant mirna bol-mir159 regulates gut microbiota composition in mice: Evidence of the crosstalk between plant mirnas and intestinal microbes. J Agr Food Chem. 2023;71:16160–73 10.1021/acs.jafc.3c06104

40. Betti F, Ladera-Carmona MJ, Weits DA et al. Exogenous mirnas induce post-transcriptional gene silencing in plants. Nat Plants. 2021;7:1379–88 10.1038/s41477-021-01005-w

41. Shahid S, Kim G, Johnson NR et al. Micrornas from the parasitic plant cuscuta campestris target host messenger rnas. Nature. 2018;553:82–85 10.1038/nature25027

42. Cai Q, Qiao L, Wang M et al. Plants send small rnas in extracellular vesicles to fungal pathogen to silence virulence genes. Science. 2018;360:1126–29 10.1126/science.aar4142

43. Wang M, Weiberg A, Dellota E et al. Botrytis small rna bc-sir37 suppresses plant defense genes by cross-kingdom rnai. Rna Biology. 2017;14:421–28 10.1080/15476286.2017.1291112

44. Jian J, Liang X. One small rna of targets and silences gene in common wheat. Microorganisms. 2019;7 10.3390/microorganisms7100425

45. Silvestri A, Fiorilli V, Miozzi L et al. In silico analysis of fungal small rna accumulation reveals putative plant mrna targets in the symbiosis between an arbuscular mycorrhizal fungus and its host plant. BMC Genomics. 2019;20:169 10.1186/s12864-019-5561-0

46. Finkel OM, Salas-González I, Castrillo G et al. A single bacterial genus maintains root growth in a complex microbiome. Nature. 2020;587:103–08 10.1038/s41586-020-2778-7

47. Penno C, Tremblay J, O’Connell Motherway M et al. Analysis of small non-coding rnas as signaling intermediates of environmentally integrated responses to abiotic stress. In: Couée I (ed.). Plant abiotic stress signaling, New York, NY: Springer US. 403–27. Retreived from 10.1007/978-1-0716-3044-0_22

48. Bolger AM, Lohse M, Usadel B. Trimmomatic: A flexible trimmer for illumina sequence data. Bioinformatics. 2014;30:2114–20 10.1093/bioinformatics/btu170

49. Parada AE, Needham DM, Fuhrman JA. Every base matters: Assessing small subunit rrna primers for marine microbiomes with mock communities, time series and global field samples. Environ Microbiol. 2016;18:1403–14 10.1111/1462-2920.13023

50. Tremblay J, Yergeau E. Systematic processing of ribosomal rna gene amplicon sequencing data. GigaScience. 2019;8:1–14 10.1093/gigascience/giz146

51. Callahan BJ, McMurdie PJ, Rosen MJ et al. Dada2: High-resolution sample inference from illumina amplicon data. Nat Methods. 2016;13:581–3 10.1038/nmeth.3869

52. Wang Q, Garrity GM, Tiedje JM et al. Naive bayesian classifier for rapid assignment of rrna sequences into the new bacterial taxonomy. Appl Environ Microbiol. 2007;73:5261–7 10.1128/AEM.00062-07

53. Quast C, Pruesse E, Yilmaz P et al. The silva ribosomal rna gene database project: Improved data processing and web-based tools. Nucleic Acids Res. 2013;41:D590–6 10.1093/nar/gks1219

54. Vilo C, Dong Q. Evaluation of the rdp classifier accuracy using 16s rrna gene variable regions. Metagenomics. 2012;1 10.4303/mg/235551

55. Kerckhof FM, Courtens EN, Geirnaert A et al. Optimized cryopreservation of mixed microbial communities for conserved functionality and diversity. PLoS One. 2014;9:e99517 10.1371/journal.pone.0099517

56. Yu B, Yang Z, Li J et al. Methylation as a crucial step in plant microrna biogenesis. Science. 2005;307:932–5 10.1126/science.1107130

57. Zhao Y, Mo B, Chen X. Mechanisms that impact microrna stability in plants. RNA Biol. 2012;9:1218–23 10.4161/rna.22034

58. Baudoin E, Benizri E, Guckert A. Impact of artificial root exudates on the bacterial community structure in bulk soil and maize rhizosphere. Soil Biology and Biochemistry. 2003;35:1183–92 10.1016/S0038-0717(03)00179-2

59. Carillo P, Gibon Y, contributors P. Protocol: Extraction and determination of proline. PrometheusWiki. 2011

60. Bramucci AR, Focardi A, Rinke C et al. Microvolume DNA extraction methods for microscale amplicon and metagenomic studies. ISME Commun. 2021;1:79 10.1038/s43705-021-00079-z

61. Love MI, Huber W, Anders S. Moderated estimation of fold change and dispersion for rna-seq data with deseq2. Genome Biology. 2014;15 10.1186/s13059-014-0550-8

62. Lin H, Peddada SD. Analysis of compositions of microbiomes with bias correction. Nat Commun. 2020;11:3514 10.1038/s41467-020-17041-7

63. Nishioka T, Elsharkawy MM, Suga H et al. Development of culture medium for the isolation of flavobacterium and chryseobacterium from rhizosphere soil. Microbes Environ. 2016;31:104–10 10.1264/jsme2.ME15144

64. Kolmogorov M, Yuan J, Lin Y et al. Assembly of long, error-prone reads using repeat graphs. Nat Biotechnol. 2019;37:540-+ 10.1038/s41587-019-0072-8

65. Dai X, Zhao PX. Psrnatarget: A plant small rna target analysis server. Nucleic Acids Res. 2011;39:W155–9 10.1093/nar/gkr319

66. John B, Enright AJ, Aravin A et al. Human microrna targets. Plos Biology. 2004;2:1862–79 10.1371/journal.pbio.0020363

67. Enright AJ, John B, Gaul U et al. Microrna targets in drosophila. Genome Biol. 2003;5:R1 10.1186/gb-2003-5-1-r1

68. Kant S, Peng M, Rothstein SJ. Genetic regulation by nla and microrna827 for maintaining nitrate-dependent phosphate homeostasis in arabidopsis. PLoS Genet. 2011;7:e1002021 10.1371/journal.pgen.1002021

69. Liang G, He H, Yu D. Identification of nitrogen starvation-responsive micrornas in arabidopsis thaliana. PLoS One. 2012;7:e48951 10.1371/journal.pone.0048951

70. Liang J, He J. Protective role of anthocyanins in plants under low nitrogen stress. Biochem Biophys Res Commun. 2018;498:946–53 10.1016/j.bbrc.2018.03.087

71. Kim Y, Cho JY, Kuk JH et al. Identification and antimicrobial activity of phenylacetic acid produced by bacillus licheniformis isolated from fermented soybean, chungkook-jang. Curr Microbiol. 2004;48:312–7 10.1007/s00284-003-4193-3

72. Musthafa KS, Sivamaruthi BS, Pandian SK et al. Quorum sensing inhibition in pseudomonas aeruginosa pao1 by antagonistic compound phenylacetic acid. Curr Microbiol. 2012;65:475–80 10.1007/s00284-012-0181-9

73. Rivera D, Mora V, Lopez G et al. New insights into indole-3-acetic acid metabolism in azospirillum brasilense. Journal of Applied Microbiology. 2018;125:1774–85 10.1111/jam.14080

74. Feng Z, Xie X, Wu P et al. Phenylalanine-mediated changes in the soil bacterial community promote nitrogen cycling and plant growth. Microbiological Research. 2023;275:127447 10.1016/j.micres.2023.127447

75. Ren B, Wang XT, Duan JB et al. Rhizobial trna-derived small rnas are signal molecules regulating plant nodulation. Science. 2019;365:919-+ 10.1126/science.aav8907

76. Ledford WC, Silvestri A, Fiorilli V et al. A journey into the world of small rnas in the arbuscular mycorrhizal symbiosis. New Phytologist. 2024;242:1534–44 10.1111/nph.19394

77. Qiao SA, Gao ZY, Roth R. A perspective on cross-kingdom rna interference in mutualistic symbioses. New Phytologist. 2023;240:68–79 10.1111/nph.19122

78. Lorenz MG, Wackernagel W. Natural genetic transformation of pseudomonas stutzeri by sand-adsorbed DNA. Arch Microbiol. 1990;154:380–5 10.1007/BF00276535

79. Nielsen KM, Bones AM, Van Elsas JD. Induced natural transformation of acinetobacter calcoaceticus in soil microcosms. Appl Environ Microbiol. 1997;63:3972–7 10.1128/aem.63.10.3972-3977.1997

80. Deng WY, Fu TW, Zhang Z et al. L-lysine potentiates aminoglycosides against acinetobacter baumannii via regulation of proton motive force and antibiotics uptake. Emerg Microbes Infec. 2020;9:639–50 10.1080/22221751.2020.1740611

81. Hong SQ, Su SP, Gao Q et al. Enhancement of β-lactam-mediated killing of gram-negative bacteria by lysine hydrochloride. Microbiology Spectrum. 2023;11 10.1128/spectrum.01198-23

82. Vinolas LC, Healey JR, Jones DL. Kinetics of soil microbial uptake of free amino acids. Biology and Fertility of Soils. 2001;33:67–74 10.1007/s003740000291

83. Sauheitl L, Glaser B, Weigelt A. Uptake of intact amino acids by plants depends on soil amino acid concentrations. Environmental and Experimental Botany. 2009;66:145–52 10.1016/j.envexpbot.2009.03.009

84. Huang W, Ma DN, Zaman F et al. Identification of the lysine and histidine transporter family in camellia sinensis and the characterizations in nitrogen utilization. Hortic Plant J. 2024;10:273–87 10.1016/j.hpj.2023.01.009

85. Guo N, Hu JQ, Yan M et al. Lysine-histidine-type transporter 1 functions in root uptake and root-to-shoot allocation of amino acids in rice. Plant Journal. 2020;103:395–411 10.1111/tpj.14742

86. Svennerstam H, Ganeteg U, Nasholm T. Root uptake of cationic amino acids by arabidopsis depends on functional expression of amino acid permease 5. New Phytol. 2008;180:620–30 10.1111/j.1469-8137.2008.02589.x

87. Svennerstam H, Jämtgård S, Ahmad I et al. Transporters in arabidopsis roots mediating uptake of amino acids at naturally occurring concentrations. New Phytologist. 2011;191:459–67 10.1111/j.1469-8137.2011.03699.x

88. Bacic A, Moody SF, Clarke AE. Structural analysis of secreted root slime from maize (zea mays l.). Plant Physiol. 1986;80:771–7 10.1104/pp.80.3.771

89. Eze MO, Amuji CF. Elucidating the significant roles of root exudates in organic pollutant biotransformation within the rhizosphere. Sci Rep. 2024;14:2359 10.1038/s41598-024-53027-x

90. Hartmann M, Zeier J. L-lysine metabolism to n-hydroxypipecolic acid: An integral immune-activating pathway in plants. Plant J. 2018;96:5–21 10.1111/tpj.14037

91. Revelles O, Espinosa-Urgel M, Molin S et al. The davdt operon of pseudomonas putida, involved in lysine catabolism, is induced in response to the pathway intermediate delta-aminovaleric acid. J Bacteriol. 2004;186:3439–46 10.1128/JB.186.11.3439-3446.2004

92. Espinosa-Urgel M, Ramos JL. Expression of a pseudomonas putida aminotransferase involved in lysine catabolism is induced in the rhizosphere. Appl Environ Microbiol. 2001;67:5219–24 10.1128/AEM.67.11.5219-5224.2001

93. Moe LA. Amino acids in the rhizosphere: From plants to microbes. Am J Bot. 2013;100:1692–705 10.3732/ajb.1300033

94. Cole BJ, Feltcher ME, Waters RJ et al. Genome-wide identification of bacterial plant colonization genes. PLoS Biol. 2017;15:e2002860 10.1371/journal.pbio.2002860

95. Pranav PS, Sivakumar R, Suvekbala V et al. Genome-wide identification of root colonization fitness genes in plant growth promoting pseudomonas asiatica employing transposon-insertion sequencing. Annals of Microbiology. 2024;74:40 10.1186/s13213-024-01784-5

