## Supplementary_Tables_Figures for "Plant miRNAs and amino acids interact to shape soil bacterial communities"

Table S1: Sequences of the single-stranded synthetic miRNAs.

| **miRNA** | **Sequences (5' to 3')** |
|---|---|
| ath-miR158a-3p | UCCCAAAUGUAGACAAAGCA |
| ath-miR158b | CCCCAAAUGUAGACAAAGCA |
| ath-miR159a | UUUGGAUUGAAGGGAGCUCUA |
| ath-miR827 | UUAGAUGACCAUCAACAAACU |
| ath-miR5642b | UCUCGCGCUUGUACGGCUUU |
| sc-ath-miR158a-3p | UGAAAACAUACAUCAGACCG |
| sc-ath-miR158b | GACAAGCUCCACAUACAGAA |
| sc-ath-miR159a | AGAGAGACCUUGGGUUGAUUU |
| sc-ath-miR827 | GAUCCAAAAAGCUAAUUUCAC |
| sc-ath-miR5642b | ACGUCGUAAAACUUCACGUG<br>UCCGGGCUUUGUACCUCUUG* |

All synthesized miRNA sequences have a 2'-OH methylation at the 3' end. *Used in the isolates vs. miRNA experiment only.

Table S2: Summary of Spearman correlations between the relative abundances of root miRNAs and bacterial taxa.

| | **ASV** | **miRNA** | **$R_S$** | ***p*-value** | |
|---|---|---|---|---|---|
| N-responding ASVs correlated to N-responding miRNAs | #14 *Massilia* | ath-miR827<br>ath-miR5642b | 0.73<br>0.60 | 0.0029<br>0.020 | *<br>* |
| | #41 *Luteimonas* | ath-miR827 | 0.70 | 0.0052 | * |
| | #478 *Massilia* | ath-miR827<br>ath-miR5642b | **-0.72**<br>**-0.53** | 0.0036<br>0.044 | *<br>* |
| Non N-responding ASVs correlated to miRNAs | #53 *Flavitalea* | ath-miR159a | 0.55 | 0.036 | * |
| | #67 *Niastella* | ath-miR158a-3p | **-0.63** | 0.014 | * |
| | # 98 *Rhizobiaceae* | ath-miR5642b | **-0.57** | 0.030 | * |
| | #111 *Ktedonobacteria* | ath-miR158a-3p<br>ath-miR827 | **-0.55**<br>0.61 | 0.038<br>0.018 | *<br>* |
| | #630 *Litorilituus* | ath-miR158a-3p<br>ath-miR827 | **-0.52**<br>0.64 | 0.047<br>0.013 | *<br>* |

Table S3: Linear regressions of the relative abundance of bacterial taxa explained by the relative abundance of root miRNAs and N treatments.

| ASV | miRNA | Adjusted $R^2$ | F-statistic | *p*-value | Significant coefficients | Relationship |
|---|---|---|---|---|---|---|
| #14 | ath-miR827 | 0.6263 | 8.822 | 0.002892 * | ns | positive |
| *Massilia* | ath-miR5642b | 0.4793 | 5.295 | 0.01672 * | Nitrogen | positive |
| #67 *Niastella* | ath-miR158a-3p | 0.4002 | 4.113 | 0.03489 * | miRNA | negative |

Linearity, normality and homoscedasticity assumptions were validated.

Table S4: Specific growth phases affected by the mix of plant miRNAs

| N Source | Growth phase affected (hours) | *p*-value | Effect of plant miRNAs |
|---|---|---|---|
| Glycine | Early Exponential (15-18) | 0.004987 | Negative |
| | Exponential (19-27) | 0.005308 | Positive |
| L-isoleucine | Exponential (39-52) | 0.003337 | Positive |
| L-leucine | Exponential (39-52) | 0.0002904 | Positive |
| L-lysine | Exponential (23-38) | 0.0001937 | Positive |
| | Stationary (39-52) | 0.02804 | Negative |
| L-phenylalanine | Exponential (20-33) | 0.002841 | Positive |
| L-proline | Exponential (0-16) | 0.002502 | Negative |
| L-valine | Stationary (22-52) | 0.007254 | Positive |
| Mix of 17 L-AA | Exponential (0-5) | 0.006503 | Positive |

Paired T-tests of the area under the curve of each growth phase were performed.

Table S5: Growth phases affected by individual miRNAs (2 µM).

| N Source | miRNA | Growth phase affected (hours) | *p*-value | Effect of plant miRNAs |
|---|---|---|---|---|
| glycine | ath-miR827 | None | N.A. | N.A. |
| | ath-miR5642b | None | N.A. | N.A. |
| L-lysine | ath-miR158a-3p | Exponential (25-37) | 0.03404 | Positive |
| | ath-miR159a | Exponential (25-37)<br>Stationary (38-40) | 0.01544<br>0.03895 | Positive |
| | ath-miR827 | Exponential (25-37) | 0.03032 | Positive |
| L-proline | ath-miR827 | None | N.A. | N.A. |
| | ath-miR5642b | None | N.A. | N.A. |
| Mix of 17 L-AA | ath-miR159a | Exponential (4-28) | 0.007735 | Positive |
| | ath-miR827 | Exponential (4-28) | 0.02033 | Positive |

Paired T-tests of the area under the curve of each growth phase were performed.

Table S6: Permutational multivariate analysis of variance of the bacterial communities (16S rRNA gene) exposed to miRNAs and cultured in media with different amino acids.

| | | Df | SumOf Sqs | $R^2$ | F | Pr(>F) |
|---|---|---|---|---|---|---|
| Exposed to the mix of 5 miRNAs | **Amino acids** | 7 | 1.0225 | 0.0857 | 1.0911 | 0.329 |
| | **miRNA treatment** | 1 | 0.5729 | 0.0480 | 4.2793 | **0.011*** |
| | **Amino acids: miRNA treatment** | 7 | 1.7629 | 0.1478 | 1.881 | **0.013*** |
| | Residual | 64 | 8.5676 | 0.7184 | | |
| | Total | 79 | 11.9259 | 1 | | |
| Exposed to single miRNAs | **miRNA treatment** | 1 | 0.01848 | 0.002015 | 0.2056 | 0.894 |
| | **miRNA family** | 3 | 0.256569 | 0.027973 | 0.9515 | 0.444 |
| | **Amino acids** | 3 | 1.308915 | 0.142705 | 4.8541 | **0.001*** |
| | **miRNA treatment: miRNA family** | 3 | 0.471203 | 0.051373 | 1.7475 | 0.099 |
| | **miRNA treatment: Amino acids** | 3 | 0.271417 | 0.029591 | 1.0065 | 0.41 |
| | **miRNA family: Amino acids** | 2 | 0.2904 | 0.031661 | 1.6154 | 0.182 |
| | **miRNA treatment: miRNA family: Amino acids** | 2 | 0.083521 | 0.009106 | 0.4646 | 0.805 |
| | Residual | 72 | 6.471647 | 0.705576 | | |
| | Total | 89 | 9.172152 | 1 | | |

Table S7: Isolates affected by miRNAs at different time points (h) of their growth phase.

| | ***Acinetobacter* (ASV#6)** | | ***Chryseobacterium* (ASV#172)** | ***Raoultella* (ASV#4)** | |
|---|---|---|---|---|---|
| | **L-phenylalanine** | **Mix 17 AA** | **Mix 17AA** | **L-Lysine** | **Mix 17 AA** |
| ath-miR158a-3p | ns | positive: 2 | ns | negative:19,23,**35-37** | ns |
| ath-miR158b | positive: 23, **28-39**, 45,47,**49-51** | ns | negative:1,3,**5-22**, **26-28**, 33,**39-41** | negative: 3, **11-15**, **17-32**, **44-51** | negative:7,10, **12-19** |
| ath-miR159a | negative: 46,51 | ns | ns | positive: 16,33,35 | ns |
| ath-miR827 | ns | positive: 0,1,3 | negative: 4,5,17,19,20,21,25 | ns | ns |
| ath-miR5642b | positive: **25-29**, **31-34**, 36 | positive: **6-8**,12,49,51 | positive: 6,7 | negative: 1,5,11,23,26 ns | ns |
| Mix of 5 miRNAs | positive:0, **5-7**, 9,11, 13, **15-19**, **21-26** | ns | positive:16,18,45, 46,51 | ns | ns |

Paired T-tests for each time point were performed.

Table S8: Growth phases of the isolates affected by plant miRNAs.

| Isolates | Culture media | miRNA | Growth phase affected (hours) | *p*-value | Effect of plant miRNA |
|---|---|---|---|---|---|
| *Raoultella* (ASV#4) | L-lysine | ath-miR158b | Exponential (0-32) | 0.001736 | Negative |
| | | | Stationary (33-52) | 0.04073 | |
| | 17 L-AA mix | ath-miR158b | Exponential (0-10) | 0.03509 | Negative |
| *Acinetobacter* (ASV#6) | 17 L-AA mix | ath-miR5642b | Exponential (0-14) | 0.02326 | Positive |
| | L-Phe | ath-miR158b | 24-45 | 0.04355 | Positive |
| | | | 46-52 | 0.03668 | |
| | | ath-miR5642b | 24-45 | 0.04480 | Positive |
| | | miRNA mix | 0-2 | 0.003491 | Positive |
| | | | 3-23 | 0.00743 | |
| *Chryseobacterium* (ASV#172) | 17 L-AA mix | ath-miR158b | Exponential (4-28) | 0.00128 | Negative |

Paired T-tests of the area under the curve of each growth phase were performed.

Table S9: Genes associated to the nitrogen cycle for each isolate.

| | ***Acinetobacter*** | ***Chryseobacterium*** | ***Raoultella*** |
|---|---|---|---|
| Nitrogen Fixation | NA | NifU | NifS |
| Nitrification | NA | NA | hydroxylamine reductase |
| Denitrification | NirB<br>NirD<br>nitrous oxide reductase family maturation protein NosD | NA | -nitrate reductase subunit alpha, beta and gamma<br>-nitrate reductase molybdenum cofactor assembly chaperone --NirB<br>-NirD<br>- anaerobic nitric oxide reductase flavorubredoxin<br>-NorR |
| Anaerobic Ammonium Oxidation (Anammox) | NA | NA | NA |
| Dissimilatory Nitrate Reduction to Ammonium (DNRA) | NA | NA | NA |
| Ammonium Assimilation | -glutamine synthetase<br>-FMN-binding glutamate synthase<br>-glutamate synthase large subunit<br>-NADP-specific glutamate dehydrogenase | -glutamine synthetase III<br>-FMN-binding -glutamate synthase<br>folylpolyglutamate synthase/dihydrofolate synthase<br>-NADP-specific glutamate dehydrogenase | -glutamine synthetase<br>-glutamate synthase large & small subunits<br>-NADP-specific glutamate dehydrogenase |
| Ammonium transport | -ammonium transporter (3x) | -ammonium transporter | -AmtB ammonium transporter |
| Nitrogen Regulation | -nitrogen regulation protein NR(I)<br>-nitrogen regulation protein NR(II)<br>-P-II family nitrogen regulator | NA | -PTS IIA-like nitrogen regulatory protein PtsN<br>-nitrogen regulatory protein P-II |
| Nitrate/Nitrite Transport | formate/nitrite transporter | NA | -NarK nitrate/nitrite MFS transporter (2x)<br>-formate/nitrite transporter<br>-nitrite transporter |

| | | | |
|---|---|---|---|
| | | | -nitrate ABC transporter permease |
| Amino Acid Transport | -branched-chain amino acid transport system II carrier protein<br>-aspartate-alanine antiporter<br>-GltP glutamate/aspartate:proton symporter<br>-ProP glycine betaine/L-proline transporter<br>-PutP sodium/proline symporter<br>-YddG aromatic amino acid DMT transporter<br>- SstT serine/threonine transporter<br>-YgaH L-valine transporter subunit | -branched-chain amino acid ABC transporter substrate-binding protein<br>-glycine betaine/L-proline ABC transporter ATP-binding protein<br>-MetIQ methionine ABC transporter | -LivKHMGF high-affinity branched-chain amino acid ABC transporter<br>-branched-chain amino acid ABC transporter permease (2x)<br>-L-methionine/branched-chain amino acid transporter<br>-branched-chain amino acid ABC transporter substrate-binding protein (2x)<br>-BrnQ branched-chain amino acid transporter carrier protein<br>-HisJPQ histidine ABC transporter permease<br>-p-aminobenzoyl-glutamate transporter<br>-GltKP glutamate/aspartate:proton symporter<br>-ProY proline-specific permease<br>-ProPVWX glycine betaine/L-proline transporter<br>-PutP sodium/proline symporter<br>-AroP aromatic amino acid transporter<br>-aromatic amino acid transport family protein<br>-YddG aromatic amino acid DMT transporter (2x)<br>- cadaverine/lysine antiporter<br>-ArgT lysine/arginine/ornithine ABC transporter substrate-binding protein<br>-lysine-specific permease<br>-phenylalanine transporter<br>- DsdX D-serine transporter<br>- SstT serine/threonine transporter<br>-HAAAP family serine/threonine permease<br>-TdcC threonine/serine transporter |

| | | | |
|---|---|---|---|
| | | | -tryptophan permease<br>-TyrP tyrosine transporter<br>-YgaH L-valine transporter subunit<br>-MetINQ methionine ABC transporter |
| General Amino Acid Transport | -amino acid ABC transporter permease (4x)<br>-amino acid ABC transporter substrate-binding protein (1x)<br>-amino acid ABC transporter ATP-binding protein (2x) | NA | -YbbAP putative ABC transporter permease subunit<br>-amino acid ABC transporter ATP-binding protein (7x)<br>-amino acid ABC transporter permease (12x)<br>-amino acid ABC transporter substrate-binding protein (2x)<br>-amino acid ABC transporter permease/ATP-binding protein (2x) |
| Oligopeptide Transport | -peptide MFS transporter<br>-SbmA peptide antibiotic transporter | -peptide MFS transporter (3x) | -OppABCF oligopeptide ABC transporter<br>-DppABCF<br>dipeptide ABC transporter<br>--peptide MFS transporter<br>-SbmA peptide antibiotic transporter |
| Molecules that increase plant AA efflux | - PhzF family phenazine biosynthesis protein (2x) | - PhzF family phenazine biosynthesis protein | - PhzF family phenazine biosynthesis (2x) |

Table S10: Predicted miRNA targets linked to amino acid transportation or nitrogen regulation.

| Isolate | miRNA | Target |
|---|---|---|
| *Acinetobacter* | -ath-miR159a | **-** contig_3_cds_pgaptmp_002450_2369<br>**putP sodium/proline symporter**<br>**-** contig_3_cds_pgaptmp_002458_2377<br>**sstT serine/threonine transporter**<br>**-** contig_3_cds_pgaptmp_003018_2936<br>**glnG nitrogen regulation protein NR(I)** |
| | -ath-miR827 | **-** contig_3_cds_pgaptmp_002388_2308<br>**AzlC family ABC transporter permease** |
| | -scramble ath-miR158a-3p | **-** contig_3_cds_pgaptmp_002304_2224<br>**amino acid ABC transporter permease** |
| | -scramble ath-miR827 | **-** contig_3_cds_pgaptmp_002975_2894<br>**brnQ branched-chain amino acid transport system II carrier protein**<br>**-** contig_3_cds_pgaptmp_003288_3206<br>**proP glycine betaine/L-proline transporter** |
| *Chryseobacterium* | None | NA |
| *Raoultella* | -ath-miR158a-3p | **-** contig_1_cds_pgaptmp_004439_4327<br>**proY proline-specific permease** |
| | -ath-miR158b | **-** contig_1_cds_pgaptmp_004439_4327<br>**proY proline-specific permease** |

| | | |
|---|---|---|
| | -ath-miR827 | -contig_1_cds_pgaptmp_003052_2958<br>**livM high-affinity branched-chain amino acid ABC transporter permease**<br>-contig_1_cds_pgaptmp_002844_2750<br>**abgT p-aminobenzoyl-glutamate transporter**<br>-contig_1_cds_pgaptmp_004439_4327<br>**proY proline-specific permease**<br>- contig_1_cds_pgaptmp_002984_2890<br>**amino acid ABC transporter ATP-binding protein**<br>- contig_1_cds_pgaptmp_001739_1676<br>**amino acid ABC transporter permease/ATP-binding protein**<br>- contig_1_cds_pgaptmp_002732_2641<br>**amino acid ABC transporter permease/ATP-binding protein** |
| | -ath-miR5642b | - contig_1_cds_pgaptmp_000221_211<br>**proP glycine betaine/L-proline transporter**<br>- contig_1_cds_pgaptmp_003523_3429<br>**yddG aromatic amino acid DMT transporter**<br>- contig_1_cds_pgaptmp_001168_1118<br>**mtr tryptophan permease** |
| | -scramble ath-miR158a-3p | - contig_1_cds_pgaptmp_001168_1118<br>**mtr tryptophan permease**<br>-contig_1_cds_pgaptmp_004488_4376<br>**amino acid ABC transporter ATP-binding protein**<br>-contig_1_cds_pgaptmp_004229_4119<br>**pheP phenylalanine transporter** |

.

Supplementary figures

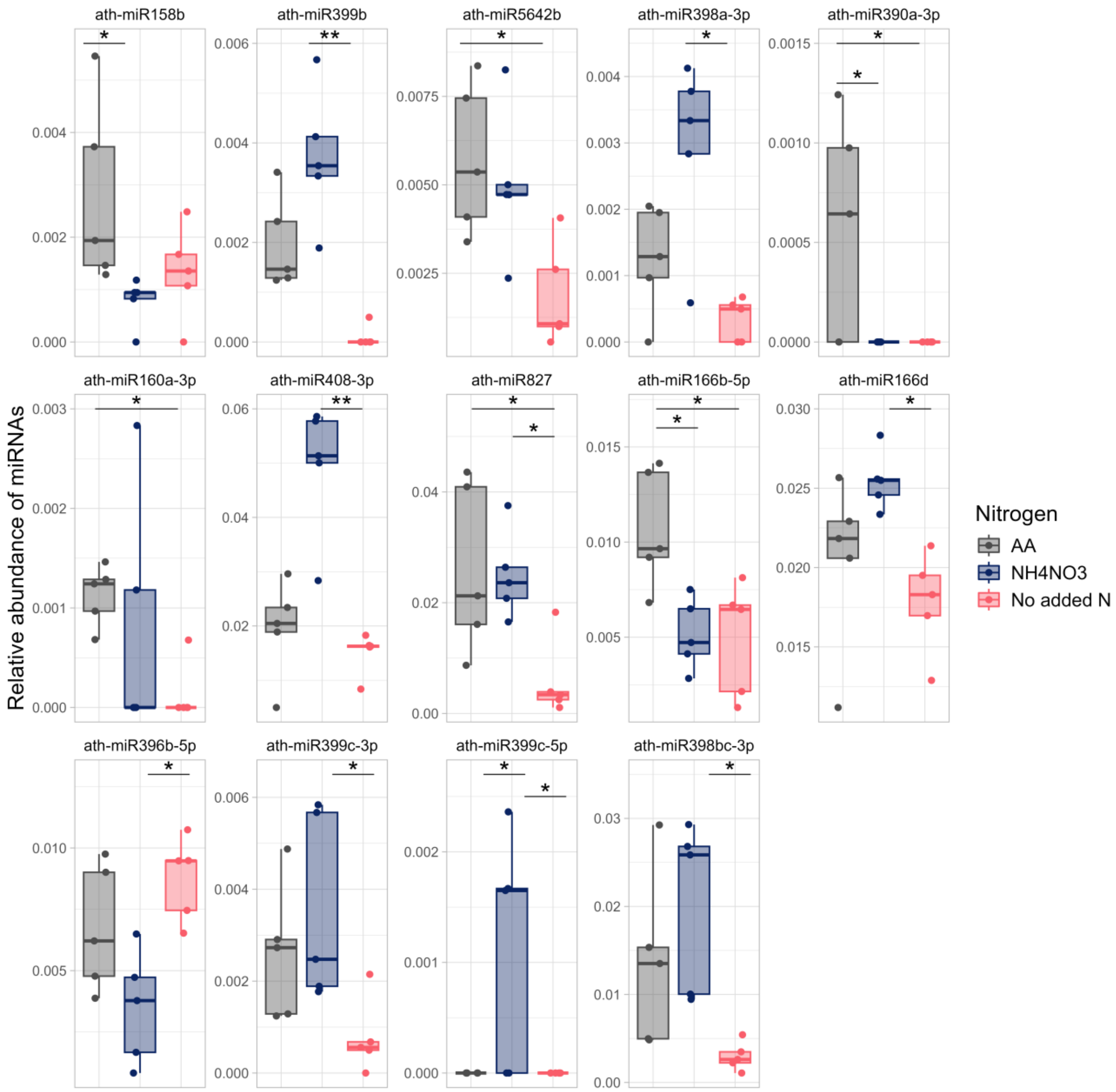


Figure S1: The relative abundance of N-responding root miRNAs. Differences in relative abundance of miRNAs between N treatments are indicated with brackets (*p*- adjusted with Holm correction <0.05).

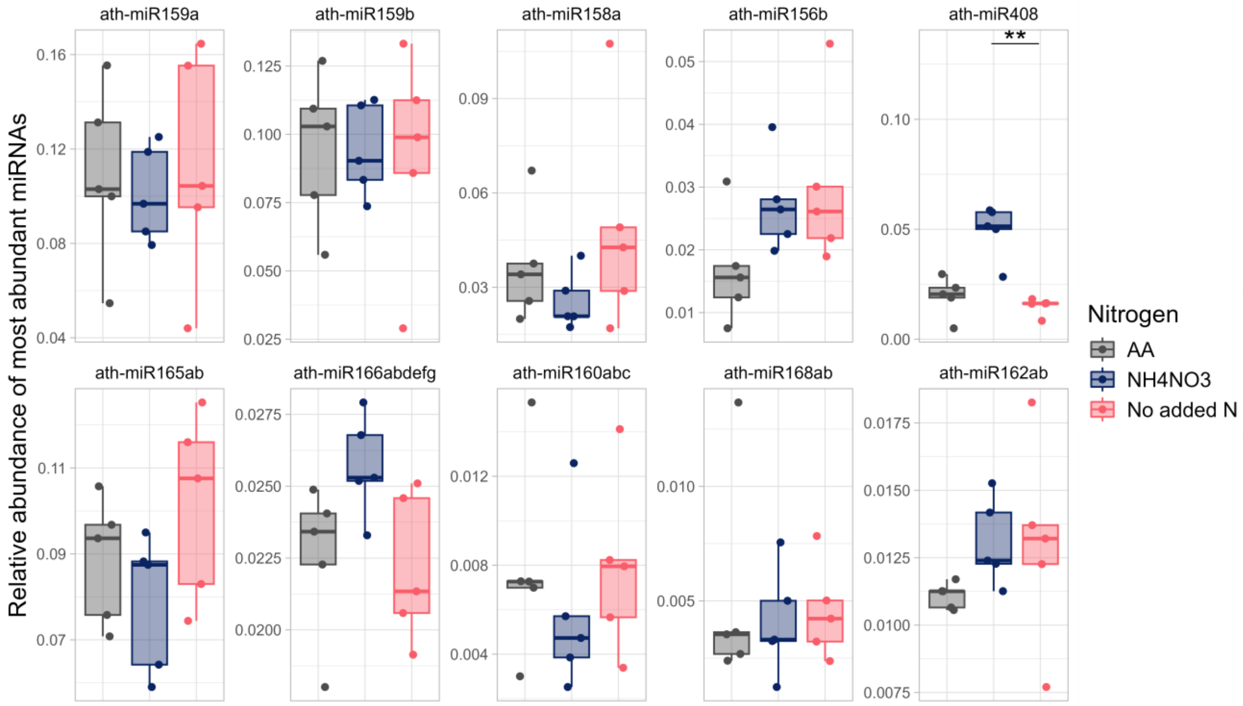


Figure S2: The relative abundance of the most abundant root miRNAs. Differences in relative abundance of miRNAs between N treatments are indicated with brackets (*p*- adjusted with Holm correction <0.05).

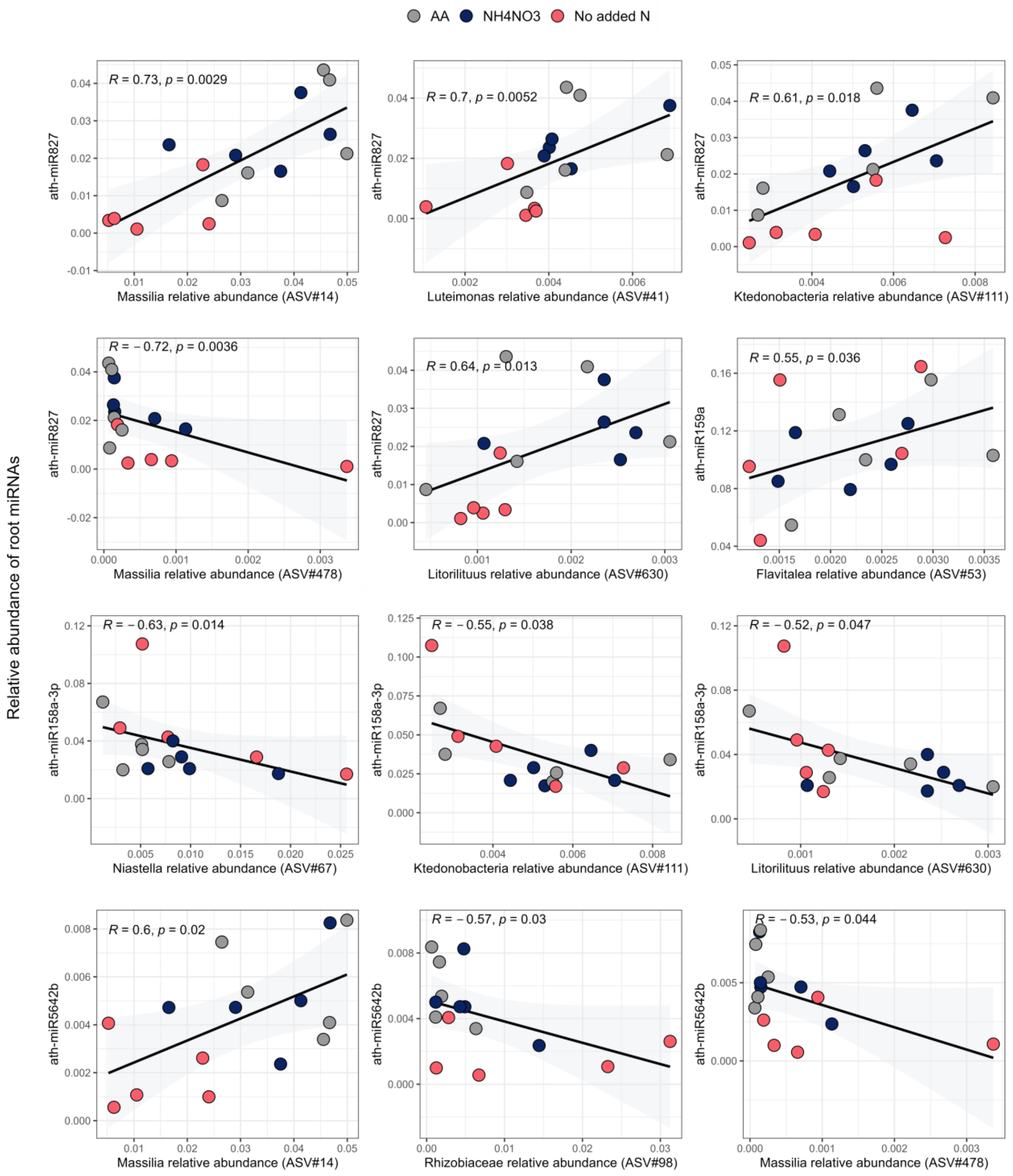


Figure S3 Spearman correlations between the relative abundance of our miRNAs of interest and bacterial ASVs (16S rRNA gene). No significant correlations were identified with ath-miR158b (*p*<0.05).

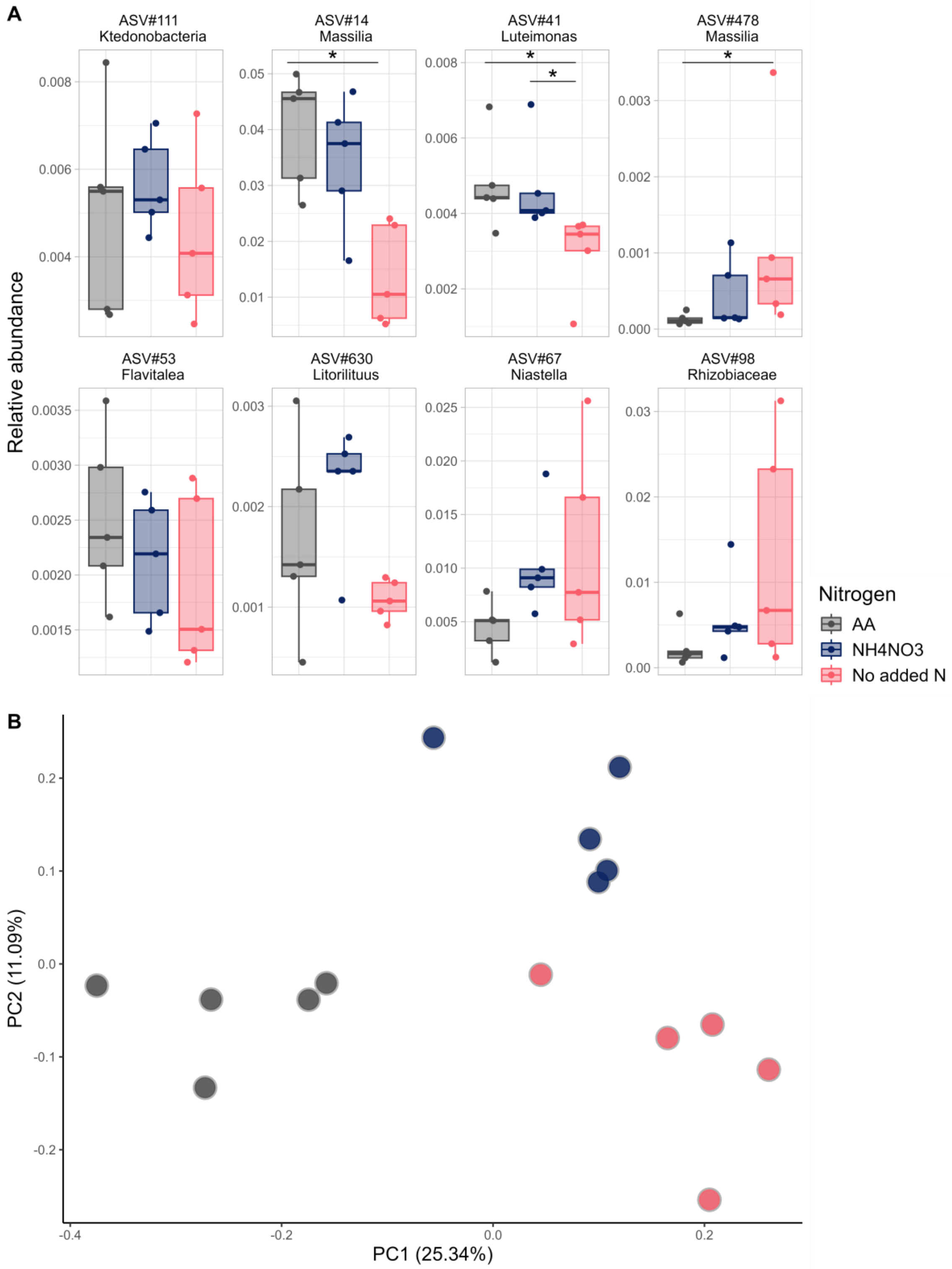


Figure S4:Bacterial taxa (16S rRNA gene) in the roots of *A. thaliana*. A. relative abundance of ASVs depending on the nitrogen treatment. Differences in relative abundance of taxa between N treatments are indicated with brackets (Dunn's test with a Benjamini-Hochberg correction $p<0.05$). B. Principle coordinate analysis that must be interpreted with the first principle component only (n=5).

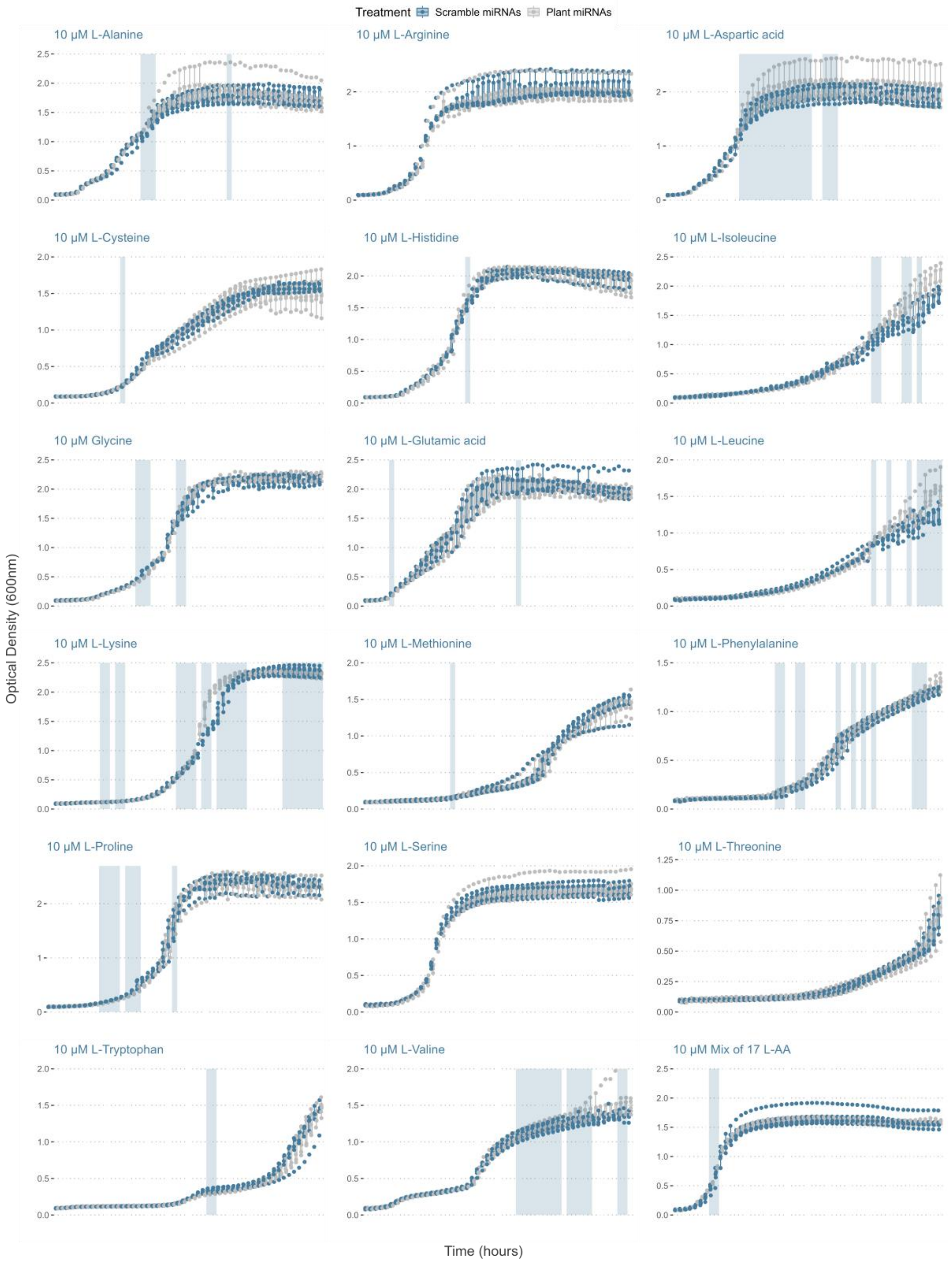


Figure S5: Growth curves, enhanced by the addition of a tetrazolium dye, of soil microbes (n=5) grown with different amino acids as a nitrogen source. The highlighted parts of the growth curve indicate differences $p<0.05$ in optical density (600 nm) of microbes treated with the mix (10 µM) plant miRNAs compared to their respective scrambled miRNAs (paired T-test).

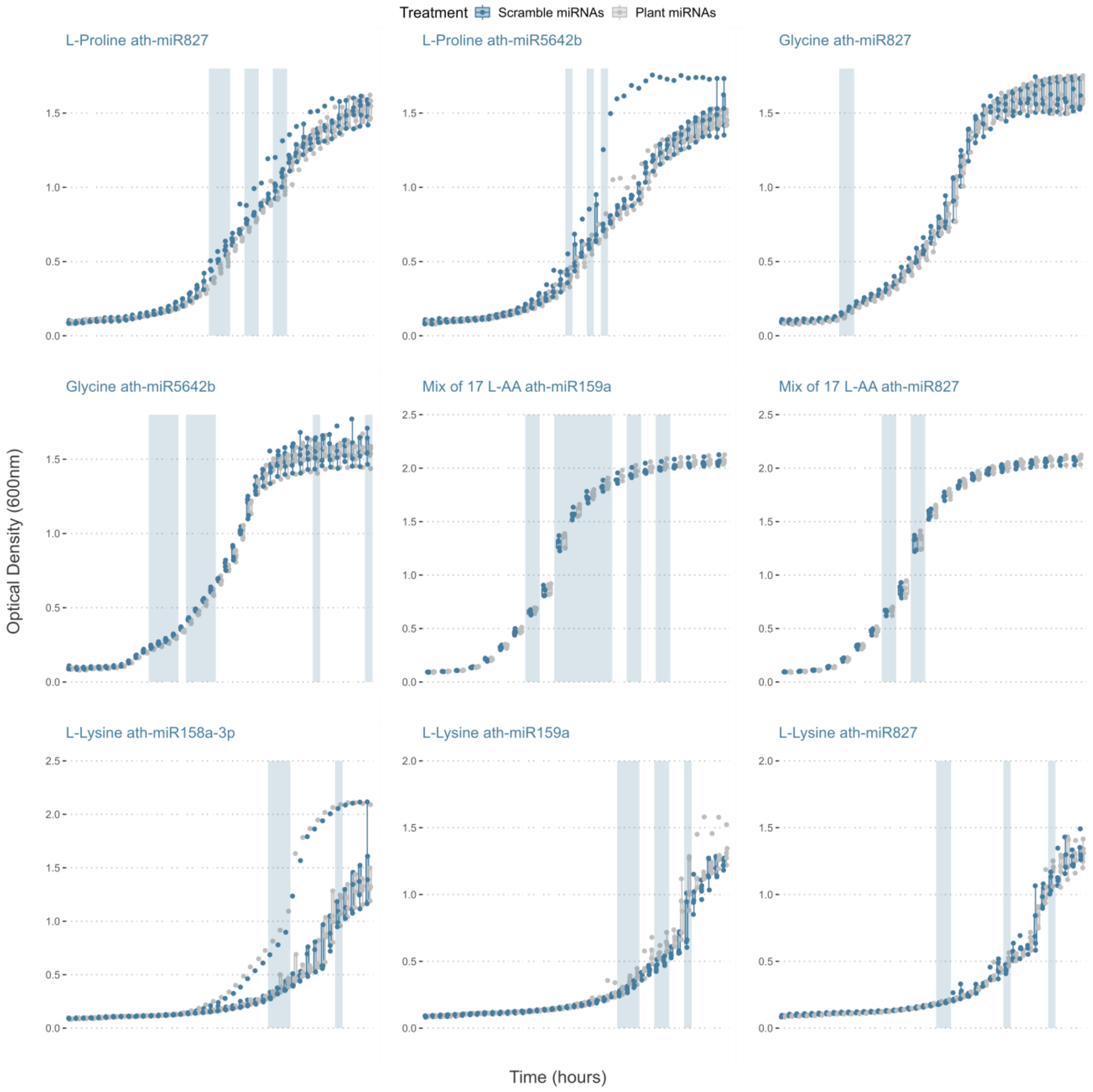


Figure S6: Growth curves, enhanced by the addition of a tetrazolium dye, of soil microbes (n=5) grown with different amino acids as a nitrogen source. The highlighted parts of the growth curve indicate differences $p<0.05$ in optical density (600 nm) of microbes treated with a single (2 µM) plant miRNA compared to its respective scrambled miRNA (paired T-test).

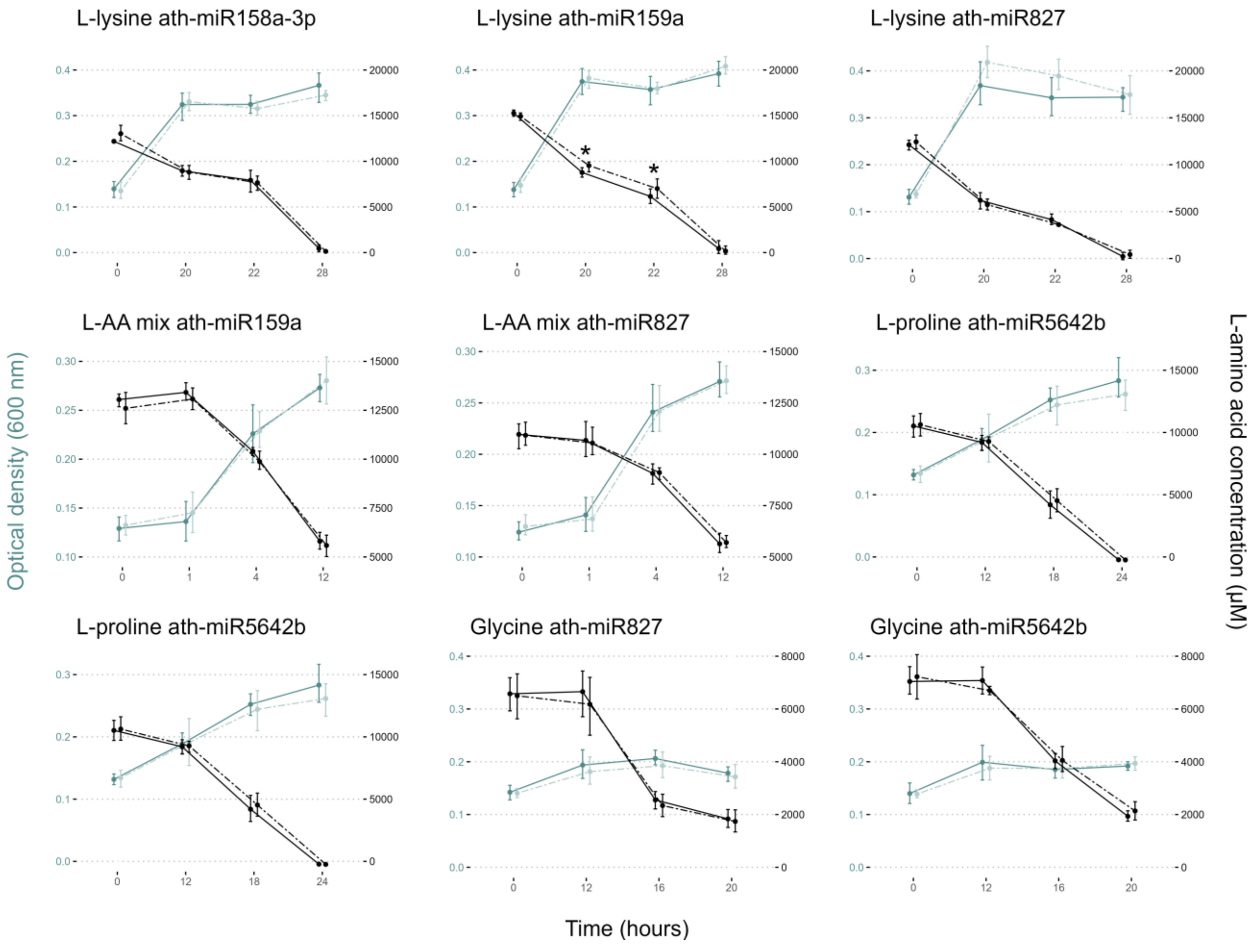


Figure S7: Microbial growth (OD600, blue lines) and amino acid consumption (black lines) over time (hours). Dashed lines indicate that the microbes were exposed to 2 µM of plant miRNAs whereas full lines indicate that the microbes were exposed to 2 µM of corresponding scrambled miRNA. Significant results are identified with a * (*p*<0.05 paired T-test) (n=5).

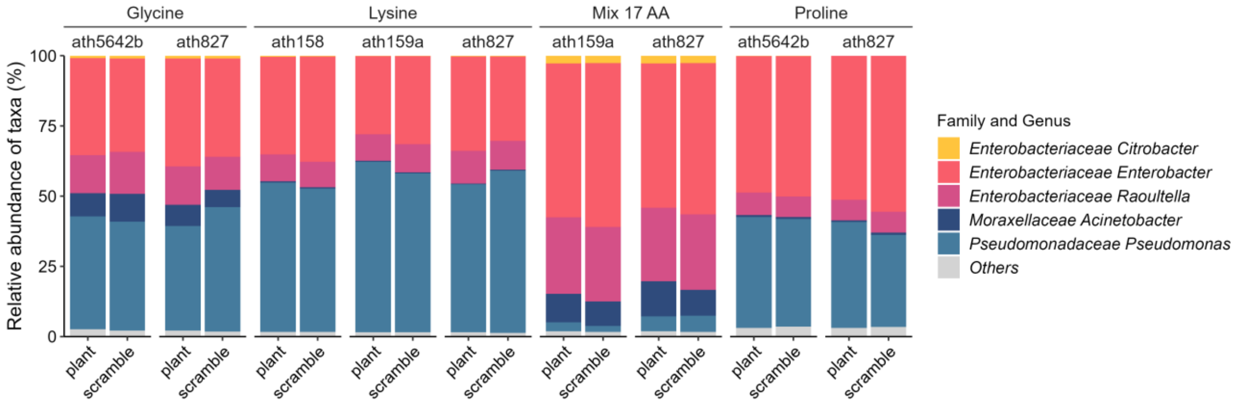


Figure S8: Mean relative abundance of bacterial taxa (16S rRNA gene) exposed to 2 µM of miRNAs (plant or scrambled) in four different amino acid sources (n=5).

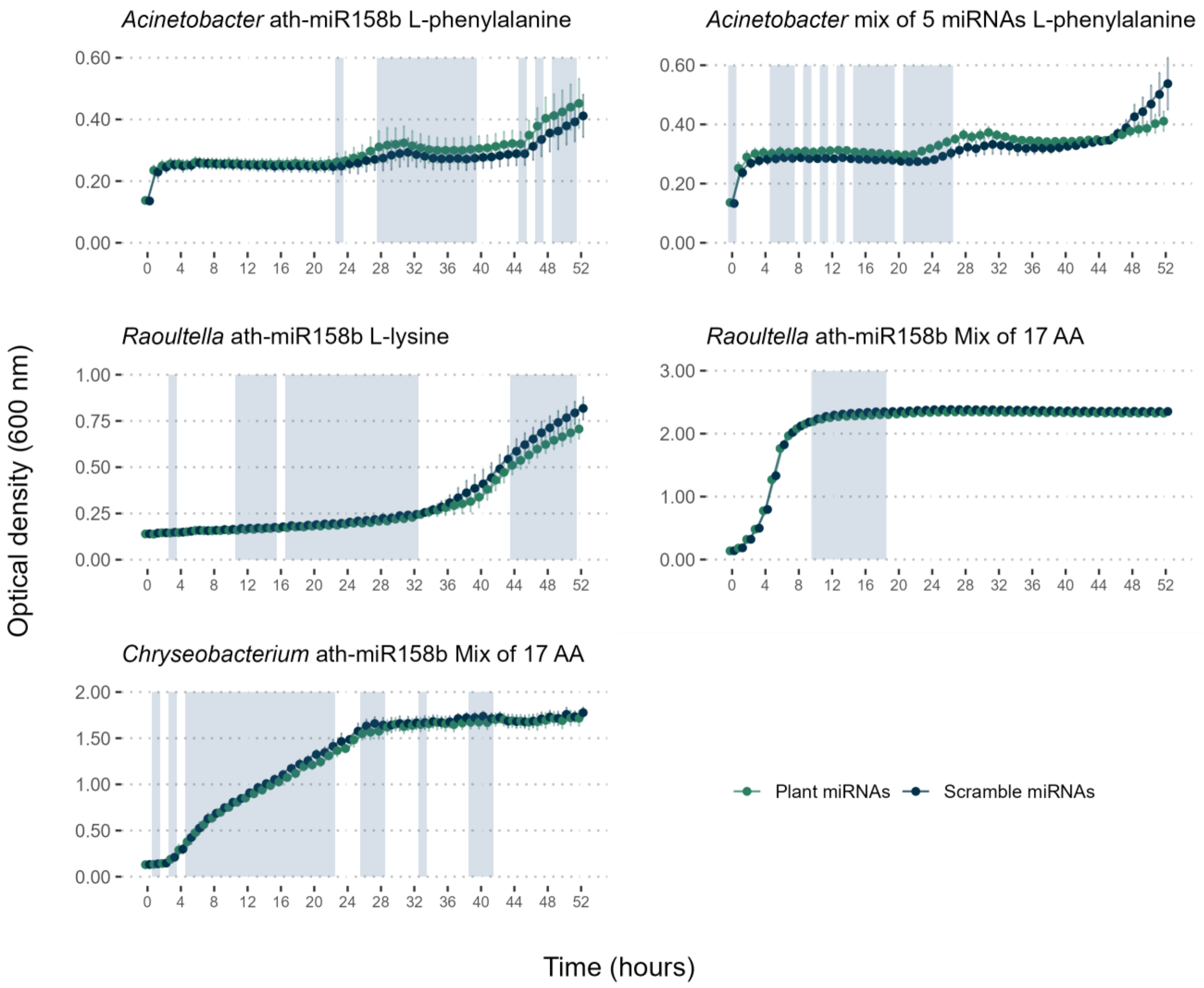


Figure S9: Growth curves of the isolates exposed to miRNAs. The highlighted parts of the growth curve indicate differences *p*<0.05 in optical density (600 nm) of isolates treated with plant miRNAs compared to the scrambled miRNA (paired T-test or paired Wilcox test) (n=5).
