## Supplementary_Methods for "Plant miRNAs and amino acids interact to shape soil bacterial communities"

**Profiles of miRNA and the bacterial community in response to fertilizer treatments *in planta.***

*Experimental design.*

*Arabidopsis thalian*a Col-0 (n=5) were grown for 21 days and under three different nitrogen treatments: a mix of 17 L-amino acids (0.190 g of N/L), a no added nitrogen control and an inorganic nitrogen control (ammonium nitrate, 0.190 g of N/L). The growth chamber was set to 12 hours of daylight, 3 hours of twilight, 3 hours of dawn, and 6 hours of darkness, with 70% humidity. During daylight, the temperature reached a maximum of 25°C, while in darkness it dropped to 20°C. Fertilizer treatments started on the fourth day after sowing and were applied every 2-3 days thereafter. On the 21st day after sowing, the roots, rhizosphere, and distant soil were sampled and flash-frozen in liquid nitrogen.

*miRNA profiling.*

The RNA was extracted from roots (RNeasy Plant Mini, Qiagen) and sent for small RNA sequencing (Illumina HiSeq4000, Centre d’expertise et de services de Génome Québec, Montreal, Canada). The small RNA sequencing reads were pre-processed and trimmed with Trimmomatic [1], filtered for quality and common contaminants were removed (bbduk). We selected reads that had a length between 18 and 27 nucleotides. These reads were mapped against a reference genome: *A. thaliana* (TAIR10/GCA_000001735.1). Alignment was carried out using the BWA parameters aln mismatch=1 and seed=5. The remaining small RNAs that mapped against the genome were compared (BLASTn) to the miRBase hairpin and miRBase mature databases using word_size=5. The BLASTn results were filtered and sequences with at least 18 bases, ≤ 3 mismatches and an overall identity percentage ≥ 85% were considered as miRNAs. As a last cleaning step, we removed all miRNAs that had been identified in our unplanted soil controls.

*Bacterial community profiling.*

The same RNA extracted from the roots, rhizosphere and bulk soil was reverse transcribed and then a PCR was performed of the V4-V5 region of the 16S rRNA gene (515F-Y: GTGYCAGCMGCCGCGGTAA and 926R: CCGYCAATTYMTTTRAGTTT [2]) for 16S amplicon sequencing on a MiSeq apparatus (Illumina) the National Research Council of Canada, (Montreal, Canada). For the taxonomic labelling steps, we treated amplicon sequencing data with the pipeline AmpliconTagger [3]. This pipeline grouped the sequences into *amplicon sequence variants* (ASVs) (100% identical sequences) with DADA2 [4] and identified their taxonomic identity with with RDP classifier using the SILVA R138 database [5-7]. The sequences that passed quality control had a mean quality greater than or equal to 25, no ambiguous bases (N) and less than 30 bases with a quality (phred Qscore) inferior to 15.

**Effects of synthetic miRNAs on a microbial community grown with different amino acids.**

*Soil microbial community*

The simplified soil community was obtained by adding 2 g of sieved (2 mm) agricultural soil from our experimental field (45.5416, -73.7173) in 20 mL of five growth media : 1) Tryptic Soy Broth, 2) Potato Dextrose Broth, 3) Minimal medium + Carbon solution + NH_4_NO_3_, 4) Minimal medium + Carbon solution + urea and 5) Minimal medium + Carbon solution + amino acids. The minimal media were supplemented with a carbon-rich solution consisting of artificial root exudates (20 mM glucose, 20 mM fructose, 12 mM sucrose, 20 mM lactic acid, 12 mM citric acid and 16 mM succinic acid) [8]. After 28 hours (200 rpm, 25 °C), the cultures were filtered (30 µm), normalized to the same optical density, then pelleted (4 °C, 15 min, 4700 *g*). The pellets were suspended in PBS, pooled, and aliquoted into sterile cryotubes. A 2x cryoprotective solution (0.6% (w/v) Tryptic Soy Broth, 10% (v/v) DMSO, and 2% (w/v) trehalose) was added to the cultures (1:1 v/v), gently mixed (5x inversion), allowed to equilibrate for 20 minutes, and stored at -80 °C [9].

*Amino acid quantification.*

We grew the simplified microbial community in four amino acid sources: L-proline, L-lysine, glycine, and the mix of 17 AA and sampled at the beginning of the experiment, at the early log-phase, the mid-log phase. For each sampling time, we removed microbial cells from the samples. They were centrifuged for 8 minutes (~6000 rpm Benchmark myFUGE^TM^ Mini) and the supernatant was transferred to a tube that was immediately placed at -80°C. The amino acids were quantified (in technical duplicates) using their respective standard curves in a colorimetric assay. For quantifying L-proline and L-lysine, we prepared a reaction mix consisting of 1% (w/v) ninhydrin, 60% (v/v) acetic acid, and 20% (v/v) ethanol [10]. Each sample (previously diluted in 70% ethanol to fit within the standard curve, final volume=50 µl) was mixed with 100 µl of the reaction mix and incubated at 95°C for 20 min. The reaction was cooled at room temperature, mixed and then 100 µl were transferred to a plate for optical density to be measured (520 nm for L-proline and 540 nm for L-lysine). To quantify glycine, the reaction mix consisted of 1% ninhydrin (w/v) diluted in 80% ethanol. The assay was carried out as previously described except that the reaction was incubated for 15 min at 75°C (520 nm). The mix of 17 AA was quantified with the colorimetric L-Amino Acid Assay Kit (CELL BIOLABS INC. MET-5054) as specified by the supplier, except for the standard curve which was our own 17-AA mix. The optical density measurements were converted to amino acid concentration (µM) using the standard curve.

**Isolates challenged with miRNAs.**

*Isolation of strains from the simplified soil community.*

We isolated the bacteria by using different solid media (MacConkey Agar, R2A Agar, Pseudomonas Isolating Agar, King’s B Agar and a Phosphate Separately autoclaved Reasoner’s 2A meant to isolate *Chryseobacterium* and *Flavobacterium* [11]. We extracted the DNA of the isolates with a physical microvolume extraction [12], precipitated the DNA (ethanol-NaCl), amplified the V4-V5 16S rRNA gene region, purified the PCR products (QIAquick PCR Purification Kit, Qiagen) and sent the samples for Sanger sequencing (Centre d’expertise et de services de Génome Québec, Montreal, Canada) (forward primer: 515FY and reverse primer:926R [2]). Consensus sequences were generated with the BioEdit Sequence Alignment Editor and compared to the 16S rRNA sequences of our previously identified ASVs with BLASTn. The isolates that had perfect matches with the responsive ASVs were further used.

*Isolates of interest exposed to miRNAs.*

We cultured the isolates with amino acids they had previously responded to in the presence of miRNAs: *Raoultella* in L-lysine and the mix of 17 AA, *Acinetobacter* in L-phenylalanine and the mix of 17 AA. *Chryseobacterium* was cultured in the mix of 17 AA rather than L-lysine, due to the isolate's inability to grow with L-lysine as a sole N source.

The amino acid use of the three isolates were also quantified as previously described. For this assay the isolates were confronted to 2 µM of individual miRNAs or 10 µM of the mix of miRNAs as well as the corresponding scrambled controls. The isolates were grown in the 17 AA mix medium and sampled at four different timepoints: beginning of the experiment, early log-phase, mid-log phase and the stationary phase (*Acinetobacter*: 0 h, 6 h, 18 h, 28 h, *Chryseobacterium*: 0 h, 8 h, 16 h, 32 h and *Raoultella* 0 h, 3 h, 6 h, 12 h). The amino acids were quantified using the colorimetric L-Amino Acid Assay Kit (CELL BIOLABS INC. MET-5054) and our own medium to generate the standard curve. We calculated the impact of plant miRNAs on each isolate’s growth and amino acid use with paired T-tests for each time point and for the area under the curve, provided the assumptions were met. Otherwise, the Wilcoxon paired test was used.

*Whole Genome sequencing of isolates.*

Isolates were cultured overnight (28 °C, 200 rpm) in M9 Medium (0.4 % m/v glucose, 0.2 % m/v casamino acids). DNA was extracted with QIAmp DNA Mini Kit (Qiagen) following the protocol for Gram negative bacteria. Quantity and quality of DNA were verified by Qubit fluorometric quantification and Nanodrop spectrophotometry prior to library preparation and Nanopore sequencing (PromethION, Oxford Nanopore technologies). The reads were combined per isolate and filtered for quality with filtlong (<https://github.com/rrwick/Filtlong>) v0.2.1 (options: --keep_percent 90 --min_length 500). The assembly was performed with Flye [13] v 2.9.3-b1797 (options  --nano-hq -i 2) and the expected genome size for each genus (option -g 3.5M for *Acinetobacter*, 5.7M for *Raoultella* (*Klebsiella*) and 5M for *Chryseobacterium*). The assembled genomes were annotated with NCBI Prokaryotic Genome Annotation Pipeline (PGAP) ([https://www.ncbi.nlm.nih.gov/refseq/annotation_prok/](https://can01.safelinks.protection.outlook.com/?url=https%3A%2F%2Fwww.ncbi.nlm.nih.gov%2Frefseq%2Fannotation_prok%2F&data=05%7C02%7CJessica.Dozois%40inrs.ca%7C96b2bc10208345721cbf08dcd8005d71%7C80c9267a177f49439958b599ca0e54c2%7C0%7C0%7C638622743282578240%7CUnknown%7CTWFpbGZsb3d8eyJWIjoiMC4wLjAwMDAiLCJQIjoiV2luMzIiLCJBTiI6Ik1haWwiLCJXVCI6Mn0%3D%7C0%7C%7C%7C&sdata=m3iqoOBhifaZnASvvKMjufpFzwQwqexrXaMClta7IPY%3D&reserved=0)).

*Putative miRNA targets within the assembled genomes.*

We used three tool to predict the targets of our plant miRNAs by mapping all 10 miRNAs (five plant sequences and five scrambled sequences) to the coding DNA sequences (CDS) of each isolate that are known to be involved in amino acid transport or general nitrogen regulation. For psRNATarget, the general parameters were used except that the extra weight in seed region was set to 0 (<https://www.zhaolab.org/psRNATarget/analysis?function=3>). For miRanda, (<https://tools4mirs.org/software/target_prediction/miranda/>) (miranda v3.3a microRNA Target Scanning Algorithm), the basic settings were used. For BLASTn, the program selected was optimized for somewhat similar sequences. The parameters were set as: Short queries: CHECK, Expect threshold = 30, Word size = 15, Max matches in a query range = 0 and Scoring Parameters: Match/Mismatch Scores = 1,-3, Gap Costs= Existence: 5 Extension 2.

*Co-culture of Arabidopsis thaliana and isolates.*

The seeds were surfaced sterilized first with 70 % ethanol, and vortexed for 2 min. The ethanol was then discarded, and 4-6 % of sodium hypochlorite (with 0.1 % of Triton X-100) was added and vortexed for 2 min. After a 5-minute wait, the sodium hypochlorite solution was discarded. The seeds were then washed with sterile water five times by shaking for 2 min in between washes. Lastly the seeds were resuspended in 1 mL of sterile water and stored at 4 °C for 2 days for cold stratification. On the day of the experiment, the seeds were inoculated with washed cultures of each isolate for 2 hours. The optical density at 600 nm (OD600) for each isolate was previously calculated to correspond to a concentration of 10^4^ CFU/µl (*Acinetobacter* (OD600=0.030), *Chryseobacterium* (OD600=0.014), and *Raoultella* (OD600=0.075)). The negative control culture medium, which was washed alongside the cultures, was used to inoculate the seeds. Ten seeds were then placed in a sterile Magenta Box containing half-strength MS (Murashige and Skoog) medium with 0.8 % agar (For 1 L: 2.2 g of Murashige and Skoog Basal Medium (M5519 Sigma), 0.25 g of MES buffer, 15 g of sucrose adjusted to a pH of 5.7 to 5.8 and 8 g of agar). The growth chamber settings were the same as in the previous *Arabidopsis* growth experiment: 12 hours of daylight, 3 hours of twilight, 3 hours of dawn, and 6 hours of darkness, with 70% humidity, with a temperature range of 20 to 25 °C. The experiment lasted for 23 days. We then determined if isolates impacted *Arabidopsis* growth by comparing different plant traits (percentage of cotyledons from 3 dpi to 6 dpi, percentage of inflorescence and fresh weight) with Kruskal-Wallis tests for each time point and for the area under the curve.
